# Senescent cells secrete chromatin components via senescence-associated extracellular particles

**DOI:** 10.64898/2025.12.12.694055

**Authors:** Sviatlana Zaretski, Jose L. Nieto Torres, Xue Lei, John P. Nolan, Malene Hansen, Peter D. Adams

## Abstract

Senescent cells influence their surroundings through the senescence-associated secretory phenotype (SASP), an assortment of secreted molecules and macromolecular complexes. Among SASP’s intracellular drivers are cytoplasmic chromatin fragments (CCFs), nuclear-derived DNA that activates the pro-inflammatory cGAS/STING pathway. While autophagy contributes to CCFs degradation, the full repertoire of CCF fates and signaling functions remains unclear. Here, we show that senescent cells release CCF components, ɣH2AX and double-stranded DNA (dsDNA), into the extracellular space via an ESCRT-independent multivesicular body pathway. Secreted CCF components localize to extracellular particles exhibiting an unusual “popcorn”-like morphology, distinct from canonical small extracellular vesicles. Notably, inhibition of autophagy enhances secretion of CCF components and particles, suggesting an inverse relationship between intracellular clearance and extracellular release. A fraction of CCF-containing extracellular particles activates cGAS-STING signaling in non-senescent proliferating cells and is enriched in the circulation of aged mice, pointing to a previously unrecognized mode of extracellular signaling by senescent cells.

## Introduction

Cellular senescence is a stable cycle arrest, often induced by various forms of molecular, cellular and tissue damage. Senescence serves a protective role by preventing the proliferation of potentially compromised, damaged, pre-malignant cells. In addition to stable proliferation arrest, senescent cells acquire a robust secretory phenotype, termed the senescence-associated secretory phenotype (SASP), which allows them to communicate with the surrounding tissue microenvironment. SASP is composed of many pro-inflammatory factors, chemokines (e.g., CCL2, CCL5, CXCL10), cytokines (e.g., IL-6, IL-8), tissue remodeling proteases (e.g., MMP-1, MMP-2)^1–3^, and Damage-Associated Molecular Patterns (DAMPs), such as DNA-binding protein HMGB1^4^, nucleic acids, and lipids. SASP also contains extracellular vesicles of different formulations, for example, small Extracellular Vesicles (sEVs), carrying diverse cargos^5^.

SASP can have beneficial effects, including promoting tissue repair and recruiting and activating immune cells to clear damaged, potentially malignant cells for the sake of tumor suppression^6,7^. Secreted HMGB1 serves as an extracellular “Alarmin” signal to neighboring cells^8,9^. However, SASP can also have detrimental effects, particularly in the context of aging and chronic disease. Persistent or excessive SASP can contribute to tissue dysfunction, chronic inflammation, and diseases of aging, including cancer^10^. SASP can also propagate senescence to neighboring cells, a phenomenon termed secondary senescence^11^. A number of SASP factors, including Notch and TGF-β family ligands, can drive this paracrine spread, potentially amplifying the negative impact of senescent cells on tissue homeostasis^12–14^. Therefore, it is of great importance to understand the full composition and mechanisms of SASP to elucidate both its protective and harmful roles as well as determine how it may function in cell-to-cell communication.

Cytoplasmic chromatin fragments (CCFs) are a recently identified specific feature of senescent cells, generated by a nucleus-to-cytoplasmic transfer of chromatin^15–18^. CCFs include heterochromatin markers, H3K9me3 and H3K27me3, but are devoid of euchromatin markers, such as H3K9ac, suggesting that CCFs are comprised predominantly of heterochromatin. CCFs are positive for the DNA damage marker ɣH2AX, but surprisingly lack its usual partner, 53BP1^18^. Mitochondria dysfunction promotes the formation of CCFs in senescent cells via a reactive oxygen species (ROS)-JNK kinase signaling pathway^16^. Intranuclear DNA damage appears to be a precursor to CCF formation, such that inhibition of MDM2, activation of p53, and enhancement of DNA repair dampen CCFs formation^19,20^. Autophagy, a conserved lysosomal degradation pathway in which ATG8/LC3B-positive autophagosomes sequester cytoplasmic cargo^21^, plays a direct role in CCFs regulation: components of the autophagy machinery have been implicated in both the formation of CCFs and their degradation^15,22^. While present in the cytoplasm, CCFs contribute to the activation of SASP through their recognition by and activation of a cytosolic nucleic acid sensor, 2’,3’-cyclic GMP-AMP synthase (cGAS)^22–25^. Activated cGAS generates the secondary messenger cyclic guanosine monophosphate-adenosine monophosphate (cGAMP) that, in turn, activates the effector protein STING, ultimately inducing the production of inflammatory cytokines and type I interferons (IFNs)^17^.

Because CCFs sit at the intersection of mitochondrial function, genome integrity, autophagy, and inflammation, and excess cytoplasmic DNA can be toxic to the cell^26,27^, defining how CCFs are generated, processed, and cleared from the cell is essential for understanding the phenotype, signaling properties, and persistence of senescent cells. In this study, we show that senescent cells secrete CCF components, including ɣH2AX and dsDNA, into the extracellular space through an ESCRT-independent multivesicular body (MVB) pathway. The CCF-related factors are released in distinctive “popcorn”-like extracellular particles, morphologically distinct from canonical sEVs, and their secretion is suppressed if autophagy is blocked. Extracellular particles are elevated in the circulation of aged mice and associate with an activity that stimulates cGAS–STING signaling in recipient non-senescent cells, pointing to a previously unknown CCF-based extracellular communication mechanism of senescent cells. These findings have broad potential implications for understanding how senescent cells manage cytoplasmic DNAs to propagate paracrine inflammation.

## Results

### Senescent cells secrete components of CCFs into the extracellular space

As one potential fate of CCFs, we wondered whether components of CCF can be secreted from senescent cells. To test for the presence of extracellular CCF components, we performed trichloroacetic acid (TCA) precipitation of proteins from conditioned serum-free media of proliferating and senescent primary human IMR90 fibroblasts (senescence induced by irradiation), as well as from senescent cells treated with MDM2 inhibitor RG7388, a previously reporeted inhibitor of CCF formation, which suppresses CCF formation nearly completely by enhancing p53-dependent DNA repair^20^ (**Figure 1A and B, Supplemental Figure 1A and B**). This analysis showed that senescent cells expelled into the extracellular media characteristic components of CCFs, ɣH2AX and H3K27me^15,18^ (**Figure 1C**). In contrast, 53BP1 and H3K9ac, both absent from CCFs^18^, were not detected in the conditioned media (**Figure 1C**). Treatment of senescent cells with MDM2 inhibitor resulted in significant suppression of secreted ɣH2AX and H3K27me3, suggesting dependence of their secretion on CCFs (**Figure 1C)**. Taken together, these results demonstrate that senescent cells secrete components of CCFs into the extracellular space.

**Figure 1:**
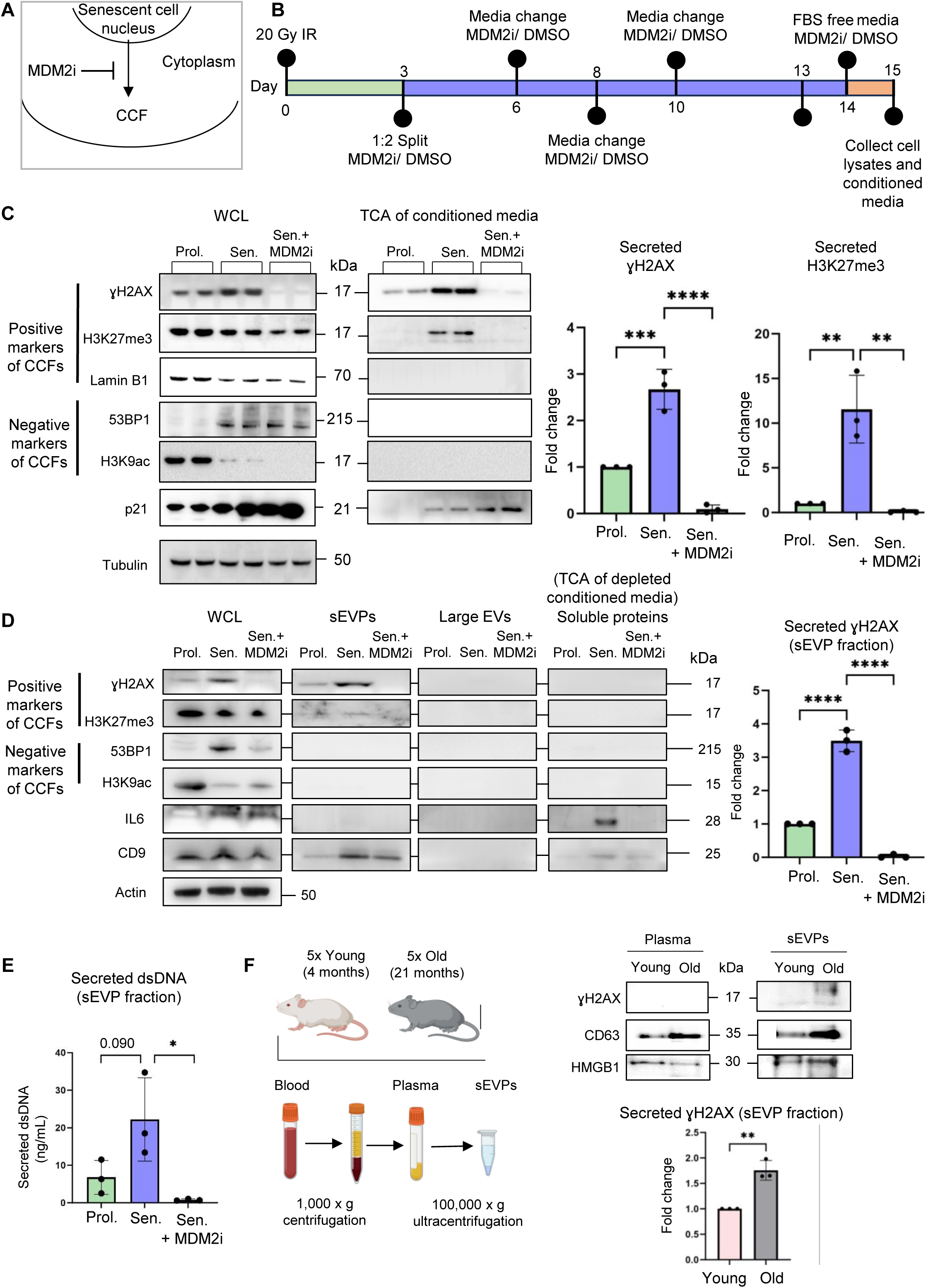
Senescent cells secrete components of CCFs into the extracellular space. (**A**) Schematic representation of a CCF exiting the nucleus into the cytoplasm of a senescent cell; this process is inhibited by the MDM2 inhibitor RG7388 (MDM2i). (**B**) Schematic of the experimental setup, timeline, and treatments, showing that cells were passaged and split 2x and treated with 100 nM MDM2i and/or DMSO vehicle on day 3 post-irradiation (20 Gy IR). Cells were dosed with fresh medium and MDM2i and/or DMSO vehicle on the indicated days. The cells were treated with FBS-free media for 24 h (orange section on the graph) before the collection. (**C**) Representative western blot of indicated proteins in whole-cell lysates (WCL) and trichloroacetic acid (TCA)–precipitated proteins from conditioned media of proliferating cells treated with DMSO vehicle or senescent IMR90 cells treated with treated with MDMi or DMSO vehicle. WCL samples were normalized to total cell number, and equal amounts of cell lysates were loaded for each condition. For TCA-precipitated samples, precipitated proteins were resuspended in identical volumes of sample buffer, and equal volumes were loaded per condition. Quantification of TCA-precipitated γH2AX and H3K27me3 was normalized to tubulin levels in the corresponding WCL and is presented as fold change relative to proliferating control cells. Data are presented as mean ± SD (n = 3 independent experiments). Statistical significance was determined using a two-sided one-way ANOVA: ***p = 0.0003, ****p < 0.0001. (**D**) Representative western blot of indicated proteins in WCL and the isolated small extracellular vesicles and particles (sEVPs), large extracellular vesicles (EV) fraction, and TCA–precipitated proteins from sEVP and large EVs-depleted conditioned media of proliferating cells treated with DMSO vehicle or senescent IMR90 cells treated with MDM2i or DMSO vehicle. WCL samples were normalized to total cell number, and equal amounts of cell lysates were loaded for each condition. For isolated sEVP, large EVs, and TCA-precipitated samples, proteins were resuspended in identical volumes of sample buffer, and equal volumes were loaded per condition. Quantification of γH2AX in the sEVP fraction was normalized to tubulin levels in the corresponding WCL and is presented as a fold change relative to proliferating control cells. Data are presented as mean ± SD (n = 3 independent experiments). Statistical significance was determined using a two-sided one-way ANOVA: ****p < 0.0001. (**E**) Quantification of secreted dsDNA in the isolated sEVP fraction from the conditioned media of proliferating cells treated with DMSO vehicle, and senescent cells treated with MDM2i or DMSO vehicle. sEVPs were isolated from equivalent numbers of cells for each condition and resuspended in identical buffer volumes prior to downstream analyses. Data are presented as mean ± SD (n = 3 independent experiments). Statistical significance was determined using a two-sided one-way ANOVA: *p=0.029. (**F**) (Left) Experimental setup outlining how blood from five young (4 months) and old (21 months) mice was collected, plasma was separated from blood cells and subsequently processed to isolate sEVPs fraction. (Right) Representative western blot of whole plasma and isolated sEVPs from five young and old mice. Equal volumes of plasma were collected from each mouse and pooled within age groups (young vs. old) prior to sEVP isolation and downstream analysis. Quantification of γH2AX in the sEVP fraction isolated from the plasma of old mice is presented as a fold change compared to γH2AX in the sEVP fraction isolated from the plasma of young mice. Data are presented as mean ± SD (n = 3 independent experiments). Statistical significance was determined using a two-sided unpaired t-test: **p=0.0025.

To further investigate the state of secreted components of CCFs, we used ultracentrifugation to separate particles and vesicles from soluble proteins based on their size and density. Multiple ultracentrifugation steps were performed to separate and isolate large (∼100-1000 nm) secreted extracellular vesicles (large EVs) and small (∼50-200 nm) secreted Extracellular Vesicles and Particles (sEVPs) from proliferating and senescent IMR90 cells, as well as senescent IMR90 cells treated with MDM2 inhibitor. By this protocol, large EVs include apoptotic bodies and plasma membrane-derived shedding vesicles formed by direct outward budding of the cell membrane^28,29^. After centrifugation, TCA precipitation was performed on the remaining depleted conditioned media to assess the efficiency of vesicle depletion and to analyze secreted soluble proteins, such as IL6 (**Figure 1D and Supplemental Figure 1C**). Using this approach, we observed increased secretion of CCF components ɣH2AX and H3K27me3 by senescent cells, specifically in the isolated fraction of CD9-containing sEVPs (**Figure 1D**). Components of CCFs were not detected in large EVs nor the fraction of soluble proteins. In addition to CCF components ɣH2AX and H3K27me^3^, we analyzed the levels of double-stranded (ds)DNA, also a component of CCFs, in the sEVP fraction^18^. We observed a similar trend of senescent cells secreting higher levels of dsDNA (**Figure 1E**). Supporting a CCF origin for these components in the sEVP fraction, the MDM2 inhibitor abolished the presence of ɣH2AX, H3K27me3, and dsDNA (**Figure 1D, E**).

The number of senescent cells increases in various tissues with age^30^. Therefore, we asked whether the CCF component ɣH2AX was more abundant in the sEVP fraction derived from plasma isolated from old mice. Indeed, analysis of the pooled plasma samples from five young (4 months old) and five old mice (21 months old) revealed an increase in ɣH2AX in the sEVP fraction from plasma of aged mice (**Figures 1F**). Together, these results indicate that senescent cells secrete cellular components characteristic of CCFs into the extracellular space in an sEVP fraction. Moreover, CCF component ɣH2AX, potentially derived from senescent cells and contained within sEVPs, circulates more abundantly in the plasma of old mice.

### Components of CCFs are secreted to the extracellular space via an ESCRT-independent pathway

The secretion of cytoplasmic components, potentially including CCFs, can be facilitated by the endosomal compartment. In this process, diverse cytoplasmic cargos, such as aggregated proteins, RNA, RNA-binding proteins, and cytoplasmic DNA, are incorporated into late endosomes, forming MVBs, through either ESCRT-dependent or ESCRT-independent mechanisms^26,31,32^. To begin to probe potential regulation of these pathways in senescent cells, we analyzed RNA-seq data derived from multiple models of senescence in IMR90 cells, including replicative senescence (RS), oncogene-induced senescence (OIS), and irradiation-induced senescence (IRS)^16,20^. This revealed a consistent upregulation of *CD9* and *CD63*, canonical markers of MVBs and sEVs^33,34^, as well as other genes implicated in secretion via the endosomal pathway (**Supplemental Figure 2A and B**). Notably, however, a subset of endosome-associated genes, predominantly those encoding components of the ESCRT machinery^35^, exhibited downregulation in all senescence models (**Supplemental Figure 2B**), suggesting a complex context-dependent remodeling of vesicular trafficking pathways during senescence. Immunofluorescent staining with both CD63 and CD9 confirmed their increased expression in puncta characteristic of MVBs in senescent cells compared to proliferating cells (**Figure 2A**). To directly test whether CCFs were recruited to MVBs, we analyzed the incorporation of CCFs into these structures (**Figure 2B**). Colocalization analysis revealed that intracellular CCFs associated with CD63, one of the markers of MVBs^36^, indicating the potential incorporation of CCFs into MVBs (**Figure 2B, C**). Together, these analyses, including comparative RNA-seq profiling across multiple senescence models and corresponding imaging-based validation, indicate an expansion of the endosomal compartment in senescent cells and demonstrate that CCFs are incorporated into MVBs, potentially on route to secretion.

**Figure 2:**
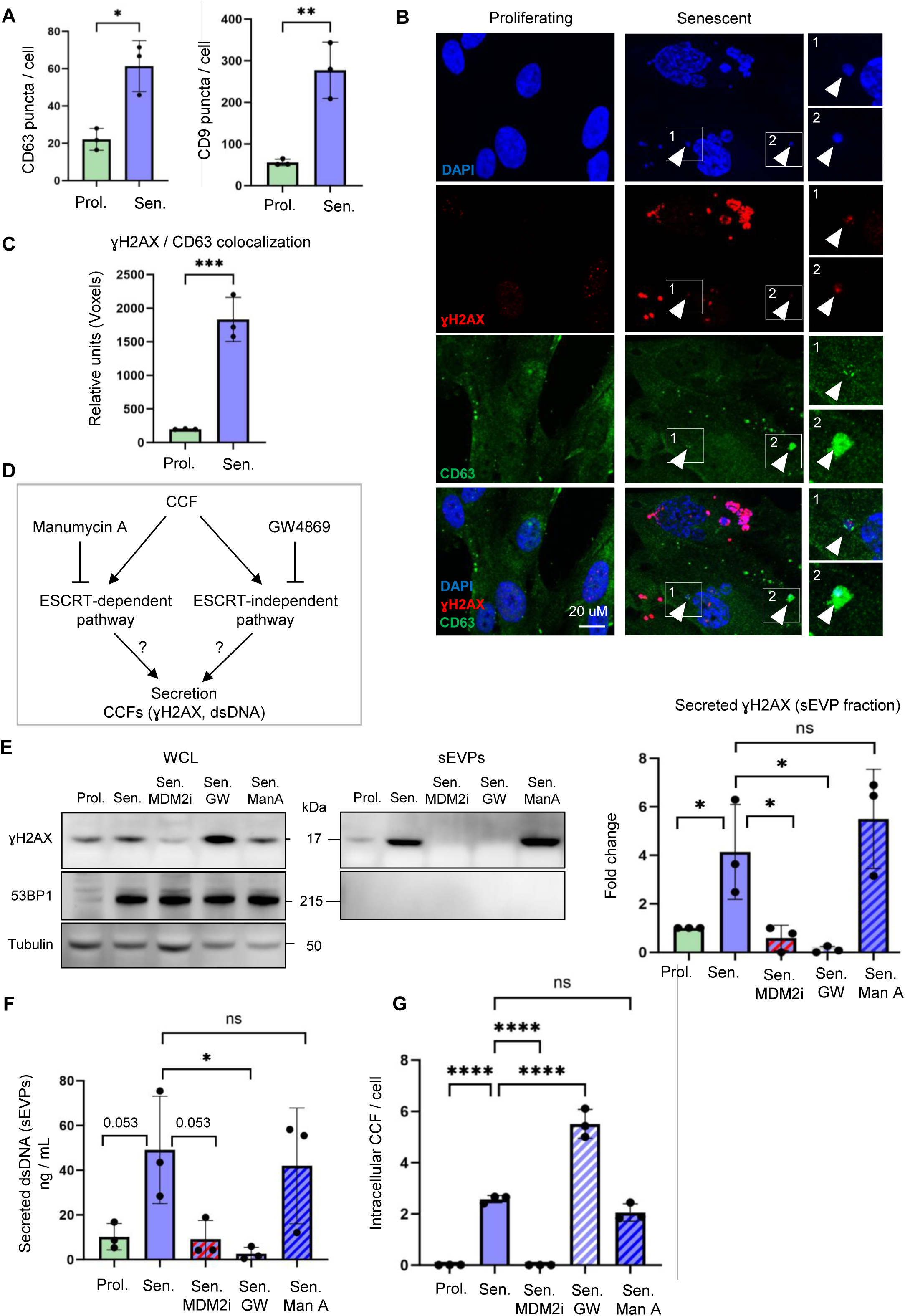
Components of CCFs are secreted to the extracellular space via an ESCRT-independent pathway. (**A**) Quantification of markers of multivesicular bodies (MVBs), CD63 and CD9 puncta/cell, in proliferating and senescent cells. The y-axis represents the total number of CD63 or CD9 punctae, normalized to the total number of nuclei. Data are presented as mean ± SD (n = 3 independent experiments). Statistical significance was determined using a two-sided unpaired t-test: *p=0.010, **p=0.0048. (**B**) Representative images and (**C**) quantifications of colocalization of cytoplasmic ɣH2AX and CD63 in proliferating and senescent cells. Numbered boxes indicate corresponding expanded regions. The y-axis represents relative units, showing the total number of colocalizing voxels in each channel, normalized to the total number of nuclei. Data are presented as mean ± SD (n = 3 independent experiments). Statistical significance was determined using a two-sided unpaired t-test: ***p = 0.0010. (**D**) Schematic representation of inhibition of ESCRT-dependent or ESCRT-independent pathways of secretion by Manumycin A or GW4869, respectively. (**E**) Representative western blot of indicated proteins in whole-cell lysates (WCL) and the isolated small extracellular vesicles and particles (sEVP) fraction from conditioned media of proliferating cells treated with DMSO vehicle, and senescent cells treated with the small-molecule inhibitor to block the formation of CCFs, RG7388 (MDM2i), or with an inhibitor of the ESCRT-independent pathway, GW4869 (GW), or an inhibitor of the ESCRT-dependent pathway, Manumycin A (ManA), or DMSO vehicle. GW4869 and ManA treatments were applied only during the final 24 h of the conditioning period. WCL samples were normalized to total cell number, and equal amounts of cell lysates were loaded for each condition. The isolated sEVP fractions were resuspended in identical volumes of sample buffer, and equal volumes were loaded per condition. Quantification of γH2AX in the sEVP fraction was normalized to tubulin levels in the corresponding WCL and is presented as a fold change relative to proliferating control cells. Data are presented as mean ± SD (n = 3 independent experiments). Statistical significance was determined using two-sided one-way ANOVA: *p=0.050 (prol. vs sen.), *p=0.044 (sen. vs sen.+MDM2i), *p=0.022 (sen. vs sen.+GW). (**F**) Quantification (Qubit) of secreted dsDNA in the isolated sEVP fraction from conditioned media of proliferating cells treated with DMSO vehicle, and senescent cells treated with MDM2i, GW4869, Manumycin A, or DMSO vehicle. GW4869 and ManA treatments were applied only during the final 24 h of the conditioning period. sEVPs were isolated from equivalent numbers of cells for each condition and resuspended in identical buffer volumes prior to downstream analyses. Data are presented as mean ± SD (n=3 independent experiments). Statistical significance was determined using two-sided one-way ANOVA: *p<0.029. (**G**) Quantification of the number of CCFs per cell in proliferating cells treated with DMSO vehicle, and senescent cells treated with GW4869, ManA, or DMSO vehicle. The y-axis represents the total number of CCF, normalized to the total number of nuclei. Data are presented as mean ± SD (n = 3 independent experiments). Statistical significance was determined using two-sided one-way ANOVA: ****p<0.0001.

To better define the role of the endosomal compartment in the secretion of CCFs, we used inhibitors to target the two main pathways of MVB biogenesis and cargo trafficking; specifically, GW4869 to inhibit nSMase required for the ESCRT-independent pathway^37^ and Manumycin A to arrest the ESCRT-dependent pathway^37^ (**Figure 2D**). Strikingly, blockade of the ESCRT-independent pathway by GW4869 suppressed secretion of ɣH2AX and dsDNA in the sEVP fraction, while inhibition of the ESCRT-dependent route by Manumycin A had no such effect (**Figure 2E-F**). Moreover, treatment of senescent cells with GW4869 led to intracellular accumulation of CCFs (**Figure 2G**), consistent with a block in trafficking out of the cell. In line with the downregulation of genes encoding the ESCRT machinery noted above (**Supplemental Figure 2B**), these data indicate that senescent cells secrete components of CCFs through MVBs via an ESCRT-independent mechanism.

### A block of autophagy upregulates the secretion of components of CCFs

Key autophagy proteins, including early-acting autophagy proteins ATG5 and ATG7, ATG8 family proteins, and autophagy receptors such as p62/SQSTM1, interact with CCFs and facilitate degradation of at least some of their components, such as Lamin B1^15,18^. Therefore, we next sought to probe the relationship between these two potential alternative fates for CCF, secretion and degradation. To inhibit degradative autophagy, we first used CRISPR/Cas9 technology to generate IMR90 cells lacking ATG16L1, a key component of the ATG8/LC3B conjugation machinery required for the early stages of autophagosome formation^38^ (**Figure 3A**). As expected, this led to a reduction in lipidated ATG8/LC3B (isoform II) protein, consistent with a block in autophagosome formation (**Supplemental Figure 3A**). We then induced these cells into senescence by irradiation and analyzed the secretion of CCFs component ɣH2AX through the sEVP fraction. Remarkably, senescent ATG16L1 knock-out cells displayed increased secretion of ɣH2AX in the sEVP fraction (**Supplemental Figure 3A**). Treatment with nSMase inhibitor GW4869 blocked this enhanced secretion in ATG16L1 knock-out cells and caused accumulation of CCFs in the cytoplasm (**Supplemental Figure 3A and B**), supporting a role for ESCRT-independent machinery and MVBs in the secretion of CCF-derived gH2AX when autophagy is blocked.

**Figure 3:**
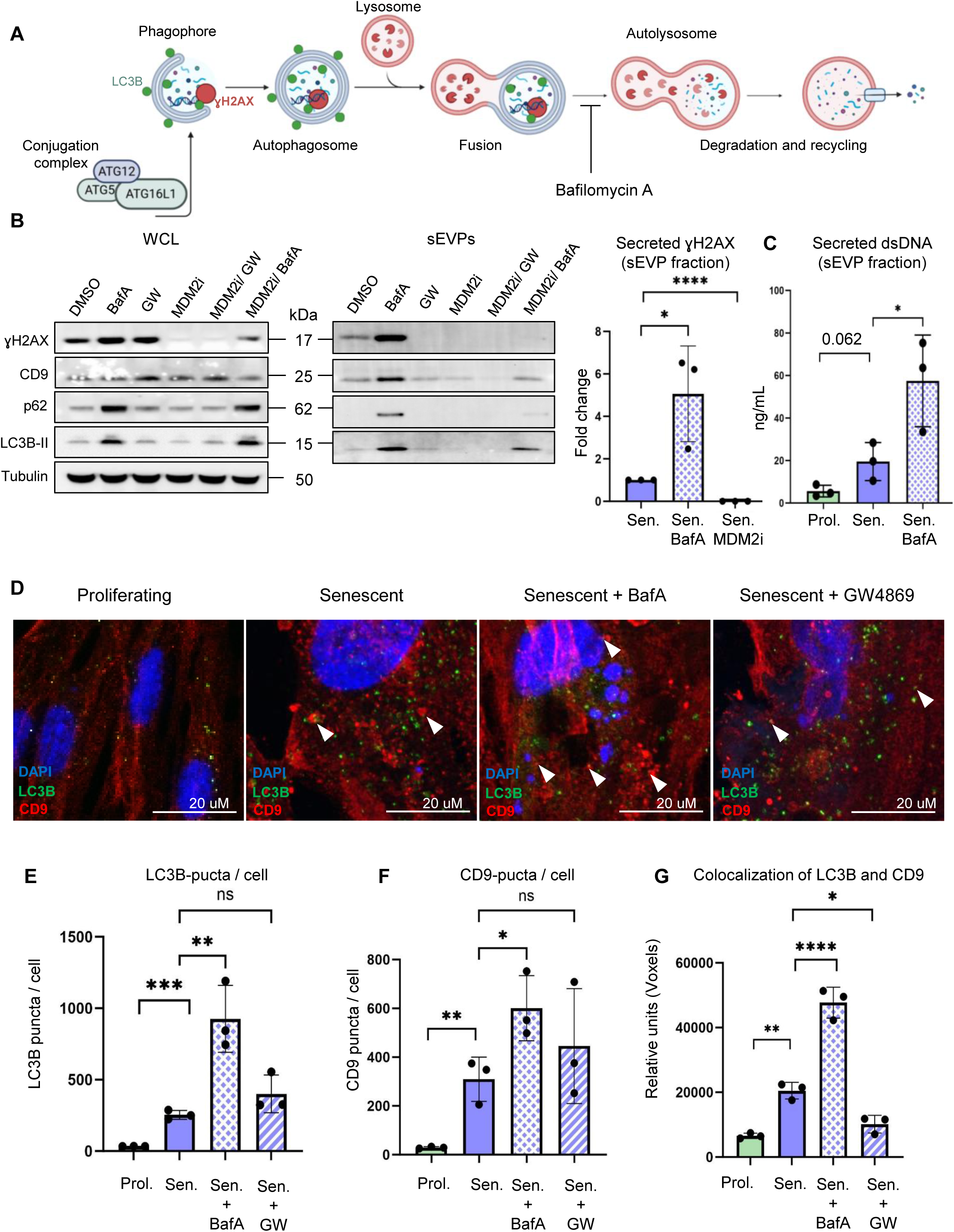
Blocking intracellular degradation of CCFs by inhibiting the process of autophagy with Bafilomycin A (BafA) upregulates secretion of the components of CCFs. (**A**) Schematic representation of the autophagy pathway, highlighting the role of ATG16L1 in conjugating ATG8/LC3B to autophagosomal membranes, and how Bafilmycin A (BafA) functions to inhibit lysosomal acidification, thus blocking degradation via autophagy. (**B**) Representative western blot of indicated proteins in whole-cell lysates (WCL) and the isolated small extracellular vesicles and particles (sEVP) fractions from conditioned media of senescent and senescent cells treated with BafA, an inhibitor of the ESCRT-independent pathway GW4869 (GW), a small-molecule inhibitor to block the formation of CCFs RG7388 (MDM2i), a combination of inhibitors, or DMSO vehicle. GW4869 and BafA treatments were applied only during the final 24 h of the conditioning period. WCL samples were normalized to total cell number, and equal amounts of cell lysates were loaded for each condition. The isolated sEVP fractions were resuspended in identical volumes of sample buffer, and equal volumes were loaded per condition. Quantification of γH2AX in the sEVP fraction was normalized to tubulin levels in the corresponding WCL and is presented as a fold change relative to senescent control cells treated with DMSO vehicle. Data are presented as mean ± SD (n=3 independent experiments). Statistical significance was determined using two-sided one-way ANOVA: ****p<0.0001, *p=0.020. (**C**) Quantification of secreted dsDNA in the isolated sEVP fraction from conditioned media of proliferating cells treated with DMSO, and senescent cells treated with BafA, or DMSO vehicle. BafA treatment was applied only during the final 24 h of the conditioning period. sEVPs were isolated from equivalent numbers of cells for each condition and resuspended in identical buffer volumes prior to downstream analyses. Data are presented as mean ± SD (n = 3 independent experiments). Statistical significance was determined using two-sided one-way ANOVA: *p<0.048. (**D**) Representative immunofluorescence images of proliferating cells treated with DMSO, and senescent cells treated with BafA, GW4869, or DMSO vehicle. GW4869 and BafA treatments were applied only during the final 24 h of the conditioning period. Arrows demonstrate colocalization events between LC3B and CD9 puncta. (**E**) Quantification of the number of LC3B-positive puncta in proliferating cells treated with DMSO, and senescent cells treated with BafA, GW4869, or DMSO vehicle. The treatments were applied only during the final 24 h of the conditioning period. The y-axis represents the total number of LC3B-positive punctae, normalized to the total number of nuclei. Data are presented as mean ± SD (n = 3 independent experiments). Statistical significance was determined using two-sided one-way ANOVA: ***p=0.0003, **p=0.013. (**F**) Quantification of the number of CD9-positive puncta CCFs per cell in proliferating cells treated with DMSO, and senescent cells treated with BafA, GW4869, or DMSO vehicle. The treatments were applied only during the final 24 h of the conditioning period. The y-axis represents the total number of CD9-positive punctae, normalized to the total number of nuclei. Data are presented as mean ± SD (n = 3 independent experiments). Statistical significance was determined using two-sided one-way ANOVA: **p=0.0060, *p=0.036. (**G**) Analysis of the colocalization of LC3B and CD9 (number of colocalizing voxels in each channel) in proliferating cells treated with DMSO, and senescent cells treated with BafA, GW4869, or DMSO vehicle. The treatments were applied only during the final 24 h of the conditioning period. The y-axis represents relative units, showing the total number of colocalizing voxels in each channel, normalized to the total number of nuclei. Data are presented as mean ± SD (n = 3 independent experiments). Statistical significance was determined using two-sided one-way ANOVA: ****p<0.0001, **p=0.0023, *p=0.014.

As an alternative approach to inhibit autophagic degradation, we employed Bafilomycin A (BafA), an inhibitor of lysosome acidification^39^ (**Figure 3A**). Like ATG16L1 KO, treatment of senescent cells with Bafilomycin A (BafA) caused a significant increase in secretion of ɣH2AX in sEVPs, as well as autophagy proteins linked to CCFs, i.e., LC3B/ATG8 and p62/SQSTM1 (**Figure 3B**). Co-treatment of senescent cells with BafA and MDM2 inhibitor showed that MDM2 inhibitor blocked secretion of ɣH2AX even in the presence of BafA, consistent with a CCF origin of ɣH2AX (**Figure 3B**). Analysis of dsDNA in the sEVP fraction showed a similar trend: dsDNA release was elevated in senescent cells and further increased by BafA (**Figure 3C**). Notably, we also observed a significant increase in the number of intracellular ATG8/LC3B- and CD9-positive puncta after BafA treatment, as well as a significant increase in the colocalization of ATG8/LC3B and CD9 (**Figure 3D-G)**. Colocalization of autophagosome marker ATG8/LC3B with endosomal marker CD9 (**Figure 3G)** can mark the formation of a hybrid organelle, the amphisome, which forms upon fusion of undegraded autophagosomes with MVBs^40,41^. In sum, these results indicate that inhibition of degradative autophagy by ATG16L1 KO or BafA enhances secretion of CCF-associated components, including ɣH2AX and dsDNA, likely through the ESCRT-independent MVB pathway.

### Senescent cells secrete a heterogeneous population of vesicles

The ultracentrifugation-based method we used to prepare the sEVP fraction efficiently recovers vesicles and particles within a similar size and density range but lacks the resolution to distinguish canonical small EVs from other secreted particles or non-canonical small vesicles^42^. Therefore, we further examined the sEVP fraction to determine its compositional heterogeneity.

First, we applied Dynamic Light Scattering (DLS) to assess vesicle/particle size and abundance. DLS analysis of the isolated sEVP fraction from proliferating, senescent, and senescent cells treated with BafA confirmed vesicles/particles within the 100–150 nm diameter range characteristic of sEVPs (**Supplemental Figure 4A-B)**. DLS also revealed increased numbers of vesicles/particles secreted by senescent cells, and this was further increased by BafA (**Supplemental Figure 4C)**.

Next, we utilized single vesicle flow cytometry (vFC) using a fluorogenic membrane probe and antibodies against EV-associated tetraspanin surface markers (i.e., CD9, CD81, CD63) to estimate vesicle/particle size, concentration, and protein/lipid composition^43,44^. vFC analysis was performed on the ultracentrifugation-isolated sEVP fractions from proliferating, senescent, and senescent cells treated with BafA. This confirmed increased secretion of canonical lipid and tetraspanin-containing sEVs from senescent cells, compared to proliferating IMR90 cells, and this population was further increased by BafA (**Supplemental Figure 4D-E**). However, this analysis also revealed another population of tetraspanin-negative, but lipid-positive vesicles/particles secreted by senescent cells, which was further increased in senescent cells treated with BafA (**Supplemental Figure 4D)**. Together, these data indicate that senescent cells release a heterogeneous population of sEVPs, which BafA treatment further amplifies.

To further resolve the heterogeneity of the ultracentrifuged fraction, we employed iodixanol density gradient centrifugation, which separates components based on their size and buoyant density (**Figure 4A**). Canonical sEVs are typically recovered within fractions of density 1.13–1.18 g/mL, whereas chromatin-associated particles and protein aggregates are known to localize to higher-density fractions (≥1.20 g/mL)^42,45,46^. By this analysis, the CCF component ɣH2AX was absent from the lower density peak canonical sEVs (CD9- and CD63-positive) fraction, but was predominantly detected in higher density fractions, containing less CD9 and CD63 (**Figure 4B**). These findings highlight the presence of potentially multiple secreted sEVP entities and suggest that ɣH2AX may not be secreted in the canonical sEV fraction.

**Figure 4:**
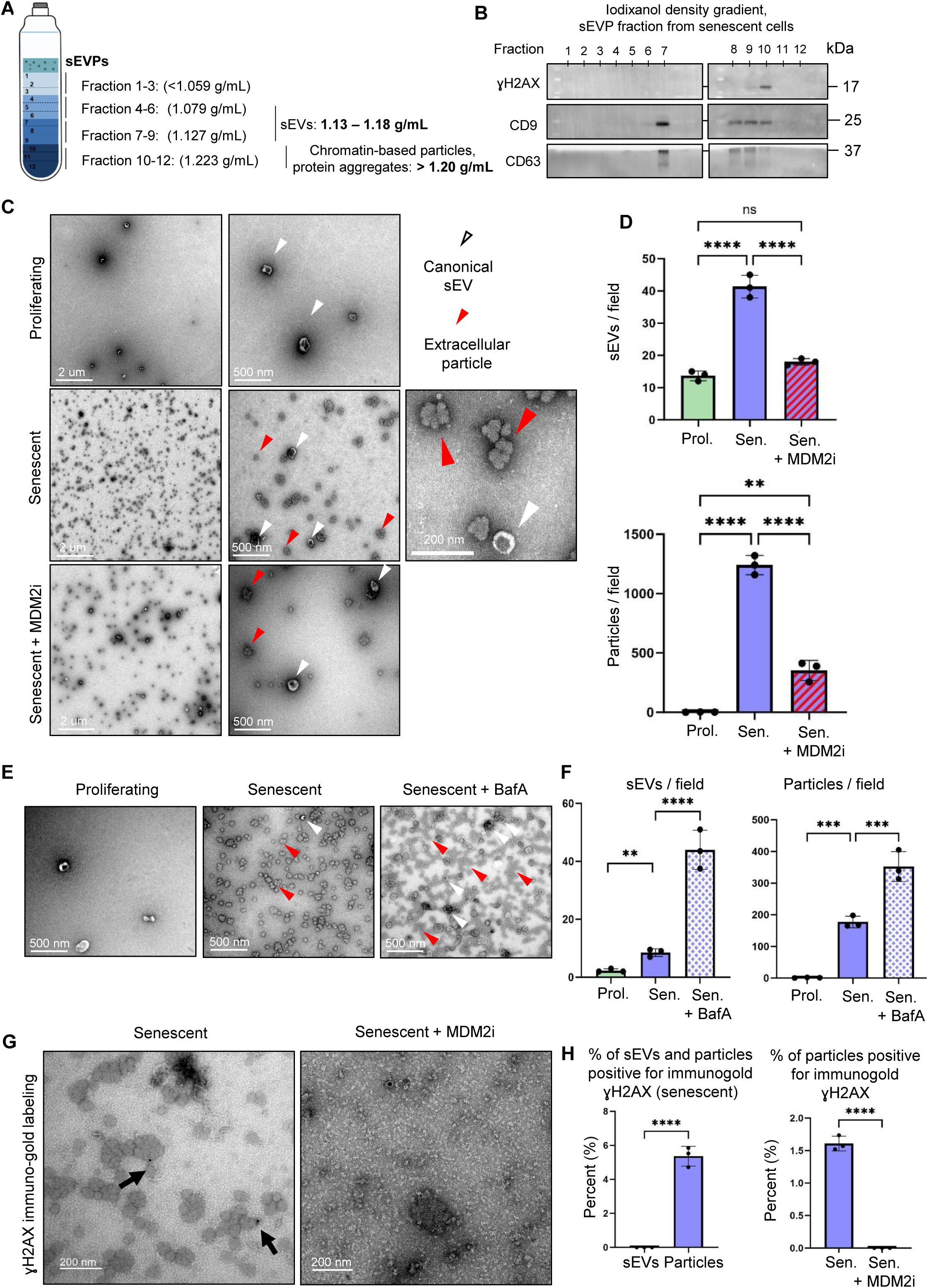
Senescent cells secrete high levels of extracellular particles as potential carriers of components of CCFs to the extracellular space. (**A**) Schematic representation of different density fractions of iodixanol density gradient and (B) representative western blot of different fractions obtained from separating the isolated small extracellular vesicles and particles (sEVP) fraction secreted by senescent cells through an iodixanol density gradient. Each fraction was analyzed for the presence of the markers of sEV, CD9 and CD63, and the component of CCFs, ɣH2AX (n = 3 independent experiments). (**C**) Transmission electron microscopy (TEM) representative images of the isolated sEVP fractions secreted by proliferating cells treated with DMSO, and senescent cells treated with the inhibitor of CCF formation, MDM2 inhibitor RG7388 (MDM2i) or DMSO vehicle. White arrows indicate canonical sEVs, and red arrows indicate extracellular particles. (**D**) Quantification of the isolated sEVs (top) and extracellular particles (bottom) in the sEVP fraction secreted by proliferating cells treated with DMSO, senescent cells treated with MDM2i, or DMSO vehicle. sEVPs were isolated from equivalent numbers of cells for each condition and resuspended in identical buffer volumes prior to downstream analyses. Data are presented as mean ± SD (n = 3 independent experiments). Statistical significance was determined using two-sided one-way ANOVA: **p=0.0017, ****p<0.0001. (**E**) TEM representative images of the isolated sEVP fractions secreted by proliferating treated with DMSO, and senescent cells treated with autophagy inhibitor Bafilomycin A (BafA), or DMSO vehicle. (**F**) Quantification of the isolated sEVs and extracellular particles secreted by proliferating cells treated with DMSO, and senescent cells treated with BafA or DMSO vehicle. sEVPs were isolated from equivalent numbers of cells for each condition and resuspended in identical buffer volumes prior to downstream analyses. Data are presented as mean ± SD (n = 3 independent experiments). Statistical significance was determined using two-sided one-way ANOVA: **p=0.0017, ***p=0.008, ****p<0.0001. (**G**) Representative TEM images of the ɣH2AX-immunogold labeling of the isolated sEVP fractions secreted by senescent cells treated with MDM2i or DMSO vehicle. sEVPs were isolated from equivalent numbers of cells for each condition and resuspended in identical buffer volumes prior to being analyzed. (**H**) Quantification of the ɣH2AX-immunogold labeling of the isolated sEVP fractions secreted by senescent cells treated with MDM2i or DMSO vehicle. **(Left)** Percentage of total small extracellular vesicles (sEVs) and extracellular particles secreted by senescent cells that stained positive for γH2AX. **(Right)** γH2AX-positive extracellular particles secreted by senescent cells treated with MDM2i or DMSO vehicle, normalized to the total number of extracellular particles per condition. Data are presented as mean ± SD (n=3 independent experiments). Statistical significance was determined using a two-sided unpaired t-test: ****p<0.0001.

### Senescent cells secrete high levels of popcorn-like extracellular particles

Given the apparent heterogeneity in the sEVP fraction, we used Transmission Electron Microscopy (TEM) for high-resolution visualization, providing critical insights into size, shape, and structural features. Of note, TEM can distinguish between spherical lipid bilayer-enclosed vesicles, such as exosomes and microvesicles, and more irregular or amorphous particles or vesicles^42^. TEM analysis of the isolated sEVP fraction confirmed the DLS and vFC analysis that, relative to proliferating cells, senescent IMR90 cells secrete an increased number of canonical sEVs, as reflected in their typical cup-shaped circular morphology and diameter of ∼100 nm (**Figure 4C-D, top panel**). Importantly, consistent with the heterogeneity detected in vFC, TEM revealed that senescent cells also secreted significant amounts of a structurally unique type of extracellular particles. Specifically, like sEVs, these extracellular particles have a diameter of ∼50-100 nm, but a very characteristic, “popcorn-like” morphology, very distinct from circular cup-shaped sEVs (**Figure 4C-D, lower panel**). Together, our TEM analysis revealed that senescent cells secrete not only increased numbers of canonical sEVs but also a distinct population of ‘popcorn-like’ extracellular particles.

The ultracentrifugation method used to precipitate and isolate the sEVP fraction could potentially create protein aggregates due to the very high applied forces. To eliminate the possibility that the observed extracellular particles were an artifact of ultracentrifugation, an alternative, more gentle method of size exclusion chromatography (SEC) was used to isolate and separate different fractions of sEVs and other particles or vesicles (**Supplemental Figure 4F**). In SEC, porous beads and differential diffusion separate components based on size: physically larger structures elute first because they cannot enter the pores, whereas smaller particles penetrate the bead matrix and therefore elute in later fractions. Using this approach, coupled with TEM, we again detected extracellular particles, specifically enriched in the later, small-particle fractions, characterized by a diameter of approximately 100-150 nm, as determined by TEM imaging (**Supplemental Figure 4F).** These results show that the extracellular particles are not specifically an artifact of ultracentrifugation. Together, the results obtained from DLS, vFC, iodixanol density gradient centrifugation, TEM, and SEC suggest that senescent IMR90 cells secrete not only canonical sEVs but also high levels of extracellular particles.

To test the conservation of these extracellular particles in other models of senescence, we tested IMR90 primary human fibroblasts induced into RS and OIS, and RS and IRS primary human melanocytes. In all cases, particles were found to be secreted; however, their secretion was more readily apparent in senescent IMR90 cells compared to senescent melanocytes (**Supplemental Figure 4G-H**). These results showed that secreted extracellular particles are a conserved qualitative feature among different models of senescence and cell types.

To extend the analysis of the extracellular particles from cell culture to *in vivo*, we investigated the presence of these particles in the plasma of young and old mice. TEM analysis of the isolated sEVP fraction from the plasma of young (4-month-old) and old mice (21-month-old) demonstrated the presence of vesicles resembling both canonical EVs and non-canonical particles secreted by senescent cells *in vitro* (**Supplemental Figure 4I**). Quantification of the isolated sEVs confirmed an increase in sEV levels in the plasma of aged mice, along with a higher abundance of extracellular particles (**Supplemental Figure 4J**). These *in vivo* findings are consistent with the hypothesis that older mice, potentially due to elevated levels of senescent cells, exhibit increased abundance of circulating sEVs and senescence-associated extracellular particles.

### Non-canonical extracellular particles carry components of CCFs into the extracellular space

Since components of CCF, sEVs, and extracellular particles were all found in the sEVP fraction, we wanted to know whether CCF components are localized to sEVs, extracellular particles, or both. To begin to address this, we first tested the impact of the genetic and pharmacologic interventions previously shown to modulate the abundance of CCF components in the sEVP fraction. The MDM2 inhibitor led to a clear reduction in the secretion of both canonical sEVs and extracellular particles (**Figure 4C-D**), in line with the decrease in secreted ɣH2AX (**Figure 1C-D**). In contrast, inhibition of autophagy, either pharmacologically with BafA or genetically through ATG16L1 knockout, resulted in a marked increase in the release of canonical sEVs and extracellular particles (**Figure 4E-F and Supplemental Figure 5A-B**), paralleling the effect on ɣH2AX secretion (**Figures 3B and Supplemental Figure 3A**). Interestingly, senescent cells treated with GW4869 to inhibit the ESCRT-independent pathway continued to secrete extracellular particles (**Supplemental Figure 5A-B**), but did not secrete ɣH2AX (**Figure 2E**). Together, these results failed to strictly link secreted CCF components to either sEVs or extracellular particles. Instead, they indicate that secretion of extracellular particles is often, but not necessarily, associated with secretion of CCFs components.

Given the complexity of interpretation of these correlative analyses, we asked directly which population, sEVs or extracellular particles, is most enriched in components of CCFs. First, TEM imaging was performed on the individual fractions isolated using the iodixanol density gradient column. Among all fractions, fraction 10 contained the highest levels of ɣH2AX (**Figure 4B**). TEM analysis revealed that this fraction was predominantly enriched in extracellular particles, whereas fractions 5 through 8 contained canonical sEVs (**Figure 4B and Supplemental Figure 5C**), indicating that a component of CCF, ɣH2AX, may associate with extracellular particles rather than canonical sEVs. We then performed immuno-gold labeling of ɣH2AX and TEM imaging of the isolated sEVs and extracellular particles. Some of the particles from senescent cells labelled positive for ɣH2AX, but this was absent from particles derived from senescent cells treated with MDM2 inhibitor (**Figure 4G-H**). Immuno-gold labeling of dsDNA as an additional marker of CCFs showed similar results (**Supplemental Figure 5D**). ɣH2AX and dsDNA label were not observed on canonical sEVs. Together, these results demonstrate that extracellular particles, but not canonical sEVs, are enriched in CCF components such as ɣH2AX and dsDNA, and therefore may serve as carriers for their extracellular secretion. To summarize, while extracellular particles and sEVs are both secreted more by senescent cells than proliferating cells, only extracellular particles detectably contain CCF components.

### The sEVP fraction is internalized by non-senescent cells and can trigger the pro-inflammatory cGAS/ STING pathway

Senescent cells influence their niche and microenvironment through paracrine or juxtacrine signaling by SASP^14,47,48^. To investigate whether the sEVP fraction might have a signaling function, we first tested for its internalization by non-senescent proliferating cells. To do this, the sEVP fraction was labelled with the fluorescent lipophilic dye PKH67, which intercalates into lipids to enable tracking of internalization by recipient cells ^49,50^. When non-senescent proliferating cells were treated with PKH67-labelled sEVPs from senescent cells, fluorescent PKH67 puncta were detected in recipient cells in proportion to the dose of PKH67 and sEVPs (**Figure 5A-C**). These data suggest that the components of the sEVP fraction can be internalized by non-senescent cells. Since senescent cell-derived extracellular particles contain dsDNA, an activator of the cGAS/STING pro-inflammatory pathway^23,24^, we next asked whether exposure of non-senescent cells to the sEVP fraction results in increased cGAMP, the product of activated cGAS and 2^nd^ messenger activator of STING. To test this, we treated proliferating IMR90 cells with the sEVP fraction from senescent cells, or, as negative controls, the same fraction from proliferating cells or senescent cells treated with the MDM2 inhibitor (**Figure 5D**; both substantially decreased in secreted dsDNA (**Figure 1E**)). We measured increased levels of cGAMP in proliferating cells treated with the sEVP fraction from senescent cells, while there was no change in cGAMP levels after treatment with the sEVP fraction from proliferating cells or senescent cells treated with the MDM2 inhibitor (**Figure 5E**). In sum, these data suggest that MDM2 inhibitor-targeted components of the senescent cell sEVP fraction, likely dsDNA, are internalized by non-senescent cells and can activate the cGAS/STING pathway.

**Figure 5:**
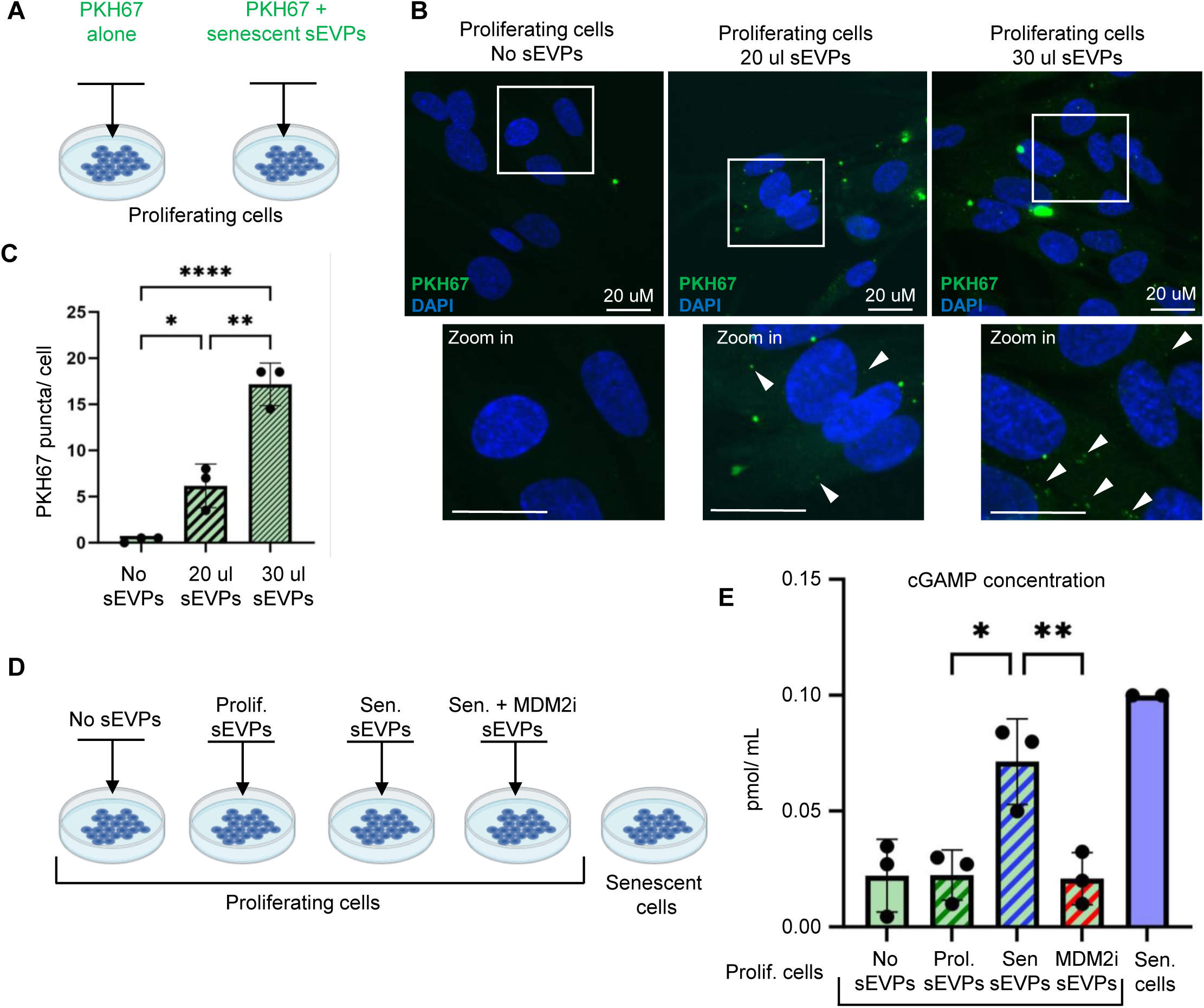
Secreted components of CCFs may be internalized by the neighboring cells and trigger the pro-inflammatory cGAS/STING pathway. (**A**) Schematic representation of the experimental approach used to test whether senescent cell–derived small extracellular vesicles and particles (EVPs) can be taken up by proliferating cells. Proliferating IMR90 cells were treated with PKH67, a green fluorescent lipophilic membrane dye, alone or with PKH67-labeled EVPs isolated from senescent cells. (**B**) Representative immunofluorescence images of proliferating cells treated with or without PKH67-labeled sEVP fractions from senescent cells, using different sEVP volumes while keeping the PKH67 lipid dye concentration unchanged. Boxes indicate expanded region and arrows indicate PKH67 puncta in recipient cells. (**C**) Quantification of internalized PKH67-labeled sEVPs by proliferating cells. Data are presented as mean ± SD (n=3 independent experiments). Statistical significance was determined using two-sided one-way ANOVA: *p=0.023, **p=0.0010, ****p<0.0001. (**D**) Schematic representation of the experimental setup: proliferating IMR90 cells were treated with no sEVPs or sEVPs secreted by proliferating cells treated with DMSO, or senescent cells treated with the inhibitor of CCF formation, MDM2 inhibitor RG7388 (MDM2i) or DMSO vehicle. (**E**) Quantification of cGAMP concentration in proliferating cells treated with no sEVPs or sEVPs (40 μL) from proliferating cells treated with DMSO, or senescent cells treated with MDM2i or DMSO vehicle. sEVPs were isolated from equivalent numbers of cells for each condition and resuspended in identical buffer volumes. Data are presented as mean ± SD (n = 3 independent experiments). Statistical significance was determined using two-sided one-way ANOVA: *p=0.011, **p=0.0094.

## Discussion

Collectively, our findings demonstrate that senescent cells secrete CCF-associated components, γH2AX, dsDNA, and H3K27me3 (but not non-CCF components 53BP1 and H3K9ac) via an ESCRT-independent MVB pathway. Secretion of γH2AX and dsDNA is antagonized by degradative autophagy. In the secreted fraction, CCF-derived components, γH2AX and dsDNA, are preferentially detected within non-canonical secretory “popcorn”-like entities, extracellular particles. The conservation of this phenomenon across multiple senescent cell types and triggers, and their detection in aged plasma, indicates extracellular particles to be a generalizable feature of senescence and potentially aging. Furthermore, we present evidence that secreted CCFs components act as signals to influence other cells.

We provide evidence that CCFs components, including γH2AX, H3K27me3 and dsDNA, are actively secreted into the extracellular space. Importantly, secretion was tightly linked to CCFs formation, as a pharmacological MDM2 inhibitor, previously shown to block CCF accumulation^20^, abrogated their extracellular release. Mechanistically, CCFs-derived cargo is secreted through an ESCRT-independent MVB pathway (see model in **Figure 6**). Interestingly, it has been previously shown that Lamin B1, another component of CCF, is degraded by autophagy. We did not detect Lamin B1 to be secreted by senescent cells, perhaps suggesting that distinct CCF components might be differentially targeted to secretory vs degradative endpoints.

**Figure 6:**
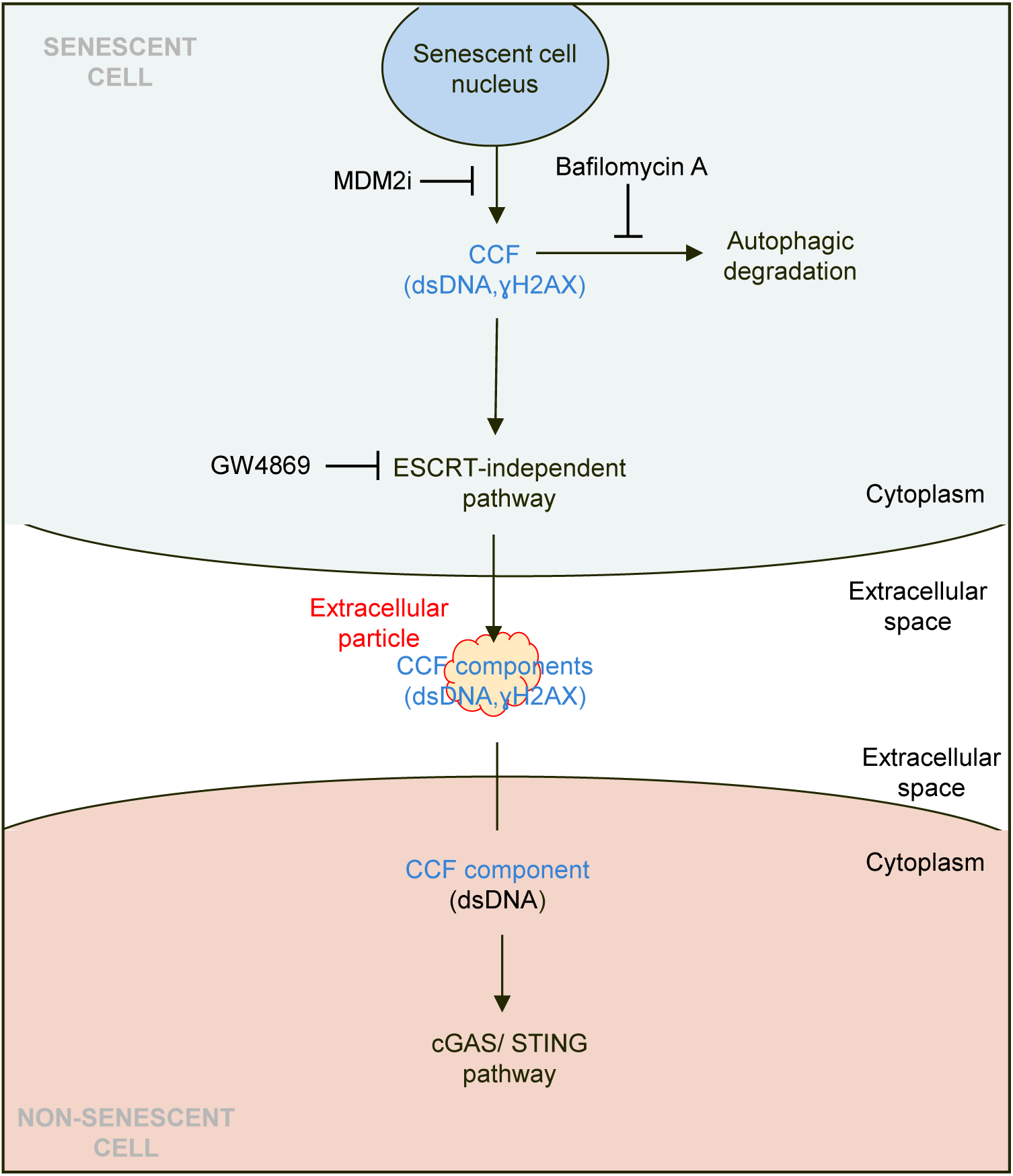
Model of the mechanism by which components of CCFs are secreted and contribute to intercellular communication. Components of CCFs are secreted from senescent cells via the endosomal compartment mediated by the ESCRT-independent pathway. These CCF-derived components are released into the extracellular space within senescence-associated extracellular particles (highlighted in red/orange). Once secreted, CCFs can be taken up by neighboring proliferating cells, where they trigger pro-inflammatory signaling responses.

These findings highlight a potential bifurcation in the cellular handling of CCFs. CCFs can be cleared by lysosomal pathways^15,51^, but when degradative autophagy is impaired so that CCFs can no longer be cleared intracellularly, CCFs are instead preferentially routed toward secretion. Consistent with this shift, inhibition of lysosomal degradation was accompanied by an accumulation of amphisomes, hybrid intermediates formed when autophagosomes fuse with MVBs for subsequent secretion^40,41^. Although the intracellular mechanisms and functional contribution of these structures to CCF handling remains to be fully defined, their appearance under conditions of blocked degradation suggests re-routing of unresolved cargo toward alternative fates. Our results suggest that cells are programmed to clear CCFs by either secretion or degradation, suggesting that their accumulation might be toxic, perhaps due to over-activation of inflammatory signals or other toxic effects of DNA in the cytoplasm^26,27^.

Immunogold labelling coupled with TEM showed that CCF components, gH2AX and dsDNA, can be detected within a subset of non-canonical “popcorn”-like entities, extracellular particles, that are secreted by senescent cells, alongside canonical sEVs. Importantly, only a relatively small fraction of these extracellular particles was positive for CCF markers, suggesting substantial heterogeneity within this particle population. Although these extracellular particles (∼50-100 nm) overlapped in size with canonical sEVs (∼100-150 nm) and were co-purified by standard ultracentrifugation, flow cytometry, density gradient fractionation, and TEM revealed them to be distinct entities. Secretion of extracellular particles was diminished upon blockade of CCF formation and enhanced by blockade of autophagy, reinforcing the regulatory link between CCF formation, autophagic flux, and the release of extracellular particles. Although the full composition and structure of extracellular particles remains to be defined, their detection across multiple senescent cell types and triggers, and also in aged plasma indicates extracellular particles to be a potential generalizable feature of senescence and aging.

Beyond their release, we present evidence that secreted CCFs exert functional effects on recipient cells. Exposure of proliferating cells to senescent cell-derived sEVPs, containing extracellular particles, triggered cGAS activation and cGAMP accumulation, providing functional evidence that extracellular CCF components, likely dsDNA, act as damage-associated molecular patterns (DAMPs), potentially capable of amplifying inflammation and propagating senescence in a paracrine manner. This suggests a direct role for extracellular CCF components in intercellular communication and systemic aging-associated inflammation. A limitation of this study is that, at this time, we cannot formally attribute the signaling function of the sEVP fraction to the extracellular particles vs canonical sEVs. However, signaling by the sEVP fraction likely depends on CCFs and extracellular particles because it was blocked by an MDM2 inhibitor, and immuno-TEM showed that CCFs components are enriched in extracellular particles compared to sEVs.

Together, these findings highlight a senescence-associated secretory pathway, revealing that CCFs can be eliminated not only through autophagic degradation but also via release in extracellular particles with probable functional cell–cell signaling properties. This work expands the current understanding of the SASP, positioning extracellular CCF components as potential mediators of inflammaging. However, important questions remain, including the precise molecular machinery governing CCF secretion as well as extracellular particle biogenesis, their composition, structure, stability and biodistribution *in vivo*, and the extent of their contribution to senescent cell signaling. Addressing these gaps will be essential for evaluating whether targeting CCFs secretion represents a viable therapeutic strategy to modulate the effects of senescent cells, for example to mitigate chronic inflammation and improve tissue homeostasis during aging.

## Acknowledgements

We thank Dr. Karl Miller, Dr. Aaron Havas, Dr. Jessica Proulx, Marcos Garcia Teneche, Adarsh Rajesh, Michael Alcatraz, and Andrew Davis for technical support with mouse sample collection. Additionally, we would like to thank Dr. Allesandra Sacco, Dr. Zhixun Dou, Dr. Nicholas Cosford, Dr. Maximiliano D’Angelo for helpful discussion, Leslie Boyd (Sanford Burnham Prebys Cell Imaging Core), Yoav Altman (Sanford Burnham Prebyc Flow Cytometry Core), Dr. Guillaume Castillon (USCD EM Core), Chun-Teng Huang (Sanford Burnham Prebys Functional Genomics and Viral Vectors Core). Parts of Figure 6 were created using Biorender.com. J.L.N.-T. is supported by a Programa Ramon y Cajal grant RYC2021-032836-I from the Spanish Ministry of Science and Innovation and a “Ayuda a la Investigación en Ciencias de la Vida y de la Materia” grant from the Fundación Ramón Areces. This work was supported by a Larry L. Hillblom Foundation network grant (2019-A-005-NET) to P.D.A. and M.H. Work in the lab of P.D.A was supported by R01 AG071861 and P30 CA030199.

## Author contributions

S.Z., J.L.N.-T., M.H., and P.D. A. designed research; S.Z. performed the experiments and data analysis, J.L.N.-T. generated the ATG16L1 knockout IMR90 cells, J. N. assisted with the vFC experiments and data analysis; X.L. analyzed RNA-seq data and generated the heatmaps shown in Supplemental Figure 2A; and S.Z. wrote the original manuscript with input from J.L.N.-T., J.N., M.H., and P.D.A.

## Data and Materials Availability

All study data are included in the article and/or supporting information.

## Online Methods

### Cell culture, treatments, and genetic manipulations

#### Cell culture and treatments

Primary IMR90 cells (PD27) were cultured in DMEM supplemented with 10% fetal bovine serum (FBS) under physiological oxygen conditions (3.5% O₂, 5% CO₂, 37 °C). For irradiation-induced senescence, IMR90 cells at ∼50% confluency (∼4.5 × 10⁶ cells per 10-cm plate) were exposed to 20 Gy X-ray irradiation (Rad Source RS-2000 Biological System). Following irradiation, cells were allowed to recover and reach confluence over 3 days, then split and replated into 15-cm dishes.

Cells were treated with the MDM2 inhibitor RG7388 (Selleckchem) at 100 nM or with DMSO vehicle control beginning after replating and maintained for 15 days post-irradiation. Media containing RG7388 or DMSO were refreshed at each passage for proliferating cells and at each media change for senescent cells.

For conditioned media collection, two 15-cm plates per condition (corresponding to a total of ∼16–20 × 10⁶ cells per condition) were used. Twenty-four h’s prior to collection, culture media were replaced with FBS-free DMEM containing DMSO or the indicated treatment, at a volume of 12.5 mL per plate. For experiments involving GW4869, Manumycin A, or Bafilomycin A, cells were treated only during this final 24-h conditioning period at the following concentrations: GW4869 (10 μM), Manumycin A (5 μM), or Bafilomycin A (50 nM).

Conditioned media were collected 15 days post-irradiation by combining 12.5 mL of media from two plates of the same condition into a single tube, yielding a total of 25 mL of conditioned media per condition. Cell lysates were collected in parallel from the same plates.

#### siRNA transfection

Primary IMR90 cells were cultured in DMEM supplemented with10% fetal bovine serum (FBS) under physiological oxygen (3.5%). One day before irradiation to induce senescence, cells were transfected with Dharmacon siGENOME 100 nM siRNA of interest in 0.8% DharmaFECT reagent according to the manufacturer’s recommendations. The next day, the cells were irradiated at 20 Gy (Rad Source RS-2000 Biological System). The media was then replaced with fresh media. 3 and 6 days after irradiation, the transfection was repeated. The cells were cultured for an additional 3 days and then collected for analysis.

#### Lentiviral Production and Transduction (for OIS experiments)

Cell Culture and Preparation: HEK293T cells were maintained in Dulbecco’s Modified Eagle Medium (DMEM) supplemented with 10% fetal bovine serum (FBS), 2 mM L-glutamine, and 100 U/mL penicillin–streptomycin. The day prior to transfection, cells were split into a 6-well plate at ∼50–60% confluence, typically by seeding one-third of a 10 cm dish per 6-well plate.

Transfection: Lentiviral vectors were transfected using Lipofectamine 2000 (Thermo Fisher). For each well: 4 µL of Lipofectamine 2000 was diluted in 150 µL Opti-MEM and incubated for 5 minutes at room temperature. DNA mixture consisting of 2.4 µg lentiviral expression plasmid (HRAS or GFP), 1 µg psPAX2 packaging plasmid, and 0.6 µg VSVG plasmid was diluted in 150 µL Opti-MEM and incubated for 5 minutes. Lipofectamine and DNA mixtures were combined and incubated at room temperature for 30 minutes. Meanwhile, 293T cells were washed with 1 mL PBS and 700 µL Opti-MEM was added. The transfection mixture (300 µL) was added dropwise to each well, and the plates were incubated for 6 h. Transfection medium was then replaced with 1 mL complete growth medium, and cells were incubated for an additional 24 h.

Viral Collection: All subsequent steps were performed under BSL-2+ conditions. Viral supernatant was collected at 24 and 48 h post-transfection, pooled, and centrifuged at 500 × g for 3 minutes to remove cell debris. Viral supernatant could either be used fresh, concentrated, or stored at – 80°C.

#### Generation of ATG16L1 IMR90 knock-out cells

CRISPR–Cas9-mediated knockout of ATG16L1 was carried out using the Synthego Lipofection Protocol for Cas9–sgRNA ribonucleoprotein (RNP) delivery, applied to low-passage IMR90 fibroblasts (PD22). Purified SpCas9 protein was complexed with a pool of three ATG16L1-targeting sgRNAs supplied by Synthego. For each transfection, 50,000 IMR90 cells were mixed with the pre-assembled RNP complex and plated into two wells of a 24-well plate: one well for continued culture and one for assessing editing efficiency by western blot. After 72 h, cells from the maintenance well were trypsinized, diluted approximately fourfold (back to ∼50,000 cells), and subjected to a second round of RNP lipofection using identical conditions. A third transfection was performed 72 h later to maximize editing efficiency. As a negative control, IMR90 cells were lipofected with Cas9 protein alone without sgRNA.

### Secretome/ Extracellular vesicle and particle isolation

#### TCA precipitation

IMR90 cells were seeded in 2 x 15 cm plates (2,000,000 cells/plate, 25 ml total volume) per condition. 24 h after seeding, cells were irradiated to induce senescence, and media containing drugs were added. 24 h before the collection of the conditioned media, the media were changed to FBS-free media in a volume of 12.5 mL per 15 cm plate. The next day, conditioned media were collected into 50 mL centrifuge tubes, combining 12.5 mL from each plate to a total of 25 mL of the collected media per condition. The collected media were subjected to a centrifugation step of 400 × g for 10 min at 4°C to pellet and remove cells. 20 mL of the supernatant were collected, leaving 5 mL, into a separate 50 mL tube. 1% (v/v) of a 2% sodium deoxycholate solution was added to each tube and incubated on ice for 30 min after mixing. Trichloroacetic acid was added to a final concentration of 7.5% (v/v) and incubated again on ice for 60 min after mixing. Proteins were precipitated by centrifugation (15,000 g for 20 min at 4°C), and the supernatants were discarded. 20 mL of 100% ice-cold (−20°C) acetone were added to the pellets, gently vortexed, and kept at −20°C for 5 min. After centrifugation (15,000 g for 5 min at 4°C), the supernatants were discarded, and 5 ml of 100% ice-cold (−20°C) acetone were added to the pellets. The pellets were vortexed again, kept at −20°C for 5 min, centrifuged (15,000 g for 5 min at 4°C), and the supernatants were discarded. The pellets were air-dried in a chemical hood for 30 min, dissolved in 200 µl of 2X SDS/ PAGE Sample Buffer, transferred to a 1.5 mL microcentrifuge tube, and kept at −80°C until use.

#### Isolation of sEVPs by ultracentrifugation

Small particulate secretome and microvesicles were isolated from conditioned media using ultracentrifugation (Beckman Coulter Optima L-100K). First, the conditioned media were collected and centrifuged at 2000 × g for 20 minutes to remove cells, large debris, and large vesicles. The resulting supernatant was subsequently centrifuged at 10,000 × g for 30 minutes to pellet microvesicles. To isolate sEVs, the supernatant was subjected to ultracentrifugation at 100,000 × g for 2.5 h. The sEV pellet was carefully resuspended and washed with filtered PBS and subjected to 100,000 × g for 2 h. The sEV pellet was collected and resuspended in filtered PBS for downstream analyses. The final pellets of both microvesicles and sEVs were either stored at −80°C or processed immediately for characterization.

#### Size-exclusion chromatography

Extracellular vesicles and particles were isolated from conditioned cell culture medium using size exclusion chromatography (SEC) with IZON qEV columns (Izon Science, Christchurch, New Zealand), following the manufacturer’s guidelines with minor modifications. Conditioned medium was first clarified by sequential centrifugation at 2,000 × g for 20 minutes to eliminate cell debris and apoptotic bodies, followed by centrifugation at 10,000 x g for 30 minutes to remove large vesicles and particles.

Prior to sample application, the qEV column was equilibrated with sterile-filtered phosphate-buffered saline (PBS; pH 7.4) according to the manufacturer’s instructions. Typically, 500 μL of clarified sample was loaded onto the column, followed by elution with PBS. Fractions of 500 μL were collected sequentially. Eluted fractions were concentrated, using 10 kDa molecular weight cut-off centrifugal filters (Amicon Ultra, MilliporeSigma) at 4,000 × g, and subsequently stored at −80 °C until use.

#### OptiPrep density gradient purification

Extracellular vesicles and particles were separated using a discontinuous iodixanol (OptiPrep; Axis-Shield) density gradient, prepared according to established protocols with slight modifications. A working stock of iodixanol (60% w/v) was diluted in 0.25 M sucrose (10 mM Tris–HCl, pH 7.4) to generate 5%, 10%, 20%, and 40% solutions. To prepare 50 mL of 40% iodixanol, 33.3 mL of the 60% stock was combined with 16.7 mL of sucrose buffer and mixed thoroughly.

Serial dilutions with equal volumes of iodixanol solution and sucrose buffer were then performed to obtain lower concentration solutions. Density gradients were assembled in 12 mL ultracentrifuge tubes by sequentially layering iodixanol solutions. First, 3 mL of 40% iodixanol was pipetted to the bottom of the tube. Next, 3 mL of 20% solution was gently layered above using an 18-gauge needle attached to a syringe, ensuring minimal disruption of the interface. Similarly, 3 mL of 10% solution was layered above, followed by 2.5 mL of 5% iodixanol solution. Finally, 500 μL of EV suspension in PBS was carefully loaded on top of the gradient. Tubes were balanced with particle-free PBS as required.

Gradients were centrifuged at 100,000 × g for 18 h at 4 °C in a swinging-bucket rotor with minimum acceleration and deceleration settings. Following centrifugation, tubes were removed carefully, and 1 mL fractions were collected sequentially from the top of the gradient using a fraction recovery system. Each fraction was transferred to a pre-labeled 1.5 mL microcentrifuge tube. Fractions of interest were transferred into fresh 12 mL ultracentrifuge tubes, diluted with 6 mL of PBS, and mixed gently. An additional 4–5 mL of PBS was added, and samples were centrifuged at 100,000 × g for 2 h at 4 °C to pellet EVs and particles and remove residual iodixanol. Supernatants were carefully decanted, and pellets were briefly tapped to remove excess liquid.

Pellets were resuspended in buffer for TEM imaging or lysed directly in 30–50 μL of lysis buffer by vortexing for 15 s, followed by incubation on a rocker for 15–20 min. Samples were then vortexed again and centrifuged briefly (500–1,000 × g for 1 min) to collect material. The resuspended pellets or collected lysates were at −80 °C or −20 °C until further use.

### Characterization of extracellular vesicles and particles

#### Nucleic acid analysis

DNA was extracted from nanoparticles using AMPure XP beads (Beckman Coulter) according to the manufacturer’s protocol. Equal volumes of nanoparticles in PBS and lysis buffer AL (QIAGEN) were mixed and incubated with Proteinase K (20 μg/ml, QIAGEN) at 56°C for 10 minutes. The mixture was then combined with one volume each of AMPure beads, isopropanol, and PEG solution (Beckman), followed by a 5-minute incubation at room temperature (RT). DNA bound to the beads was separated from the solution using a magnet for 5 min at RT. The supernatant was removed, and bead-bound DNA was washed twice with 80% ethanol, air-dried for 5 minutes, and eluted with nuclease-free water. DNA concentration was quantified using the QuBit assay (Life Technologies). DNA extraction was performed for two independent biological replicates of each sample.

#### Dynamic light scattering analysis

The hydrodynamic size distribution of secreted extracellular particles was determined using dynamic light scattering. Particle preparations were first isolated from cell culture supernatants via differential ultracentrifugation and resuspended in sterile filtered phosphate-buffered saline (PBS). Samples were equilibrated at 25 °C and analyzed using a Zetasizer Nano ZS (Malvern Panalytical) equipped with a 633 nm laser. Data were analyzed using instrument software to determine the mean particle size and polydispersity index (PDI). Care was taken to avoid bubbles and aggregates in the cuvette to ensure accurate size determination. Performed at UCSD, Materials Research Science & Engineering Center.

#### Flow cytometry analysis of particles

Vesicle concentration, size, and tetraspanin (TS)-positive fraction were determined by single vesicle flow cytometry^43,44^ using a commercial kit (vFC Assay kit, Cellarcus Biosciences, San Diego, CA) and flow cytometer (Cytek Aurora, SBP Flow Cytometry core). Briefly, samples were stained with the fluorogenic membrane stain vFRed and a mix of fluorescent antibodies targeting CD9, CD63, and CD81, molecules often associated with some EVs, for 1h at RT. Data were analyzed using FCS Express (De Novo Software) and included calibration using vesicle size and fluorescence intensity standards. The analysis included a pre-stain dilution series to determine the optimal initial sample dilution and multiple positive and negative controls, per guidelines of the International Society for Extracellular Vesicles (ISEV)^52,53^.

#### PKH-67 labeling of particulate secretome

sEVs were fluorescently labeled with PKH67 (MINI67-1KT, Merck) following the manufacturer’s guidelines with modifications to optimize specificity. Briefly, sEVP fractions were isolated from senescent IMR90 cells as described above, with the final pellet resuspended in 100 µl of diluent C instead of PBS. As a critical control, FBS-free DMEM was incubated at 37°C for 48 h, processed in parallel through the exosome isolation workflow, and resuspended in 100 µl of diluent C. This control accounts for the potential aggregation of PKH67 in the absence of vesicles, which can otherwise confound downstream assays. For labeling, 6 µl (1×) or 12 µl (2×) of PKH67 dye was diluted in diluent C to a final volume of 50 µl. Each dye preparation was mixed with either sEV or FBS-free DMEM suspensions and incubated for 5 min at room temperature. Following labeling, 18 ml of PBS was added, and samples were centrifuged at 100,000 × g for 1.5 h to pellet PKH67-labeled sEVPs or labeled FBS-free DMEM-derived background particles. Pellets were resuspended in PBS and wrapped in foil to protect from light. Labelled samples were added directly to the proliferating IMR90 cells and incubated for 6 h. The cells were fixed with 4% PFA and imaged for analysis.

#### Transmission electron microscopy (TEM)

The isolated sEVP fraction (10 μL per condition) was loaded onto a glow-discharged copper EM grid coated with a thin carbon film. The sEVPs were allowed to adsorb to the grid for 5-10 minutes. Following adsorption, the grid was washed with distilled water and further stained with a filtered uranyl acetate solution for 1 minute. The grid was then blotted and air-dried before imaging using a Transmission Electron Microscope, Jeol 1400 plus at UCSD, Electron Microscopy Core. For each condition and independent experimental repeat, an average of 3–6 fields of view were imaged and analyzed.

#### Immuno-TEM

The isolated sEVP fraction was loaded onto a glow-discharged copper EM grid coated with a thin carbon film. The sEVPs were allowed to adsorb to the grid for 10 minutes. Following adsorption, to block nonspecific binding, the grids were incubated with 1% bovine serum albumin (BSA) in PBS for 30 min at room temperature. Additionally, the grids were incubated with or without 0.01% Triton X to permeabilize any vesicles present in the samples. Primary antibodies (1:20 dilution in 0.1% BSA, PBS) were applied to the grids for 1 h at room temperature. Following five washes in 0.1% BSA, PBS, grids were incubated with secondary antibody conjugated to 12 nm colloidal gold particles ( dilution in 0.1% BSA, PBS) for 1 h at room temperature. Grids were then washed five times with 0.1% BSA, PBS, followed by two washes with distilled water. Grids were further stained with a filtered uranyl acetate solution for 1 minute. The grid was then blotted and air-dried before imaging using a Transmission Electron Microscope, Jeol 1400 plus at UCSD, Electron Microscopy Core. For each condition and independent experimental repeat, an average of 3–6 fields of view were imaged and analyzed.

### Cellular and molecular analyses

#### cGAMP assay

Intracellular 2’3’-cyclic GMP-AMP (cGAMP) levels were quantified using the 2’3’-cGAMP ELISA Kit (Invitrogen) according to the manufacturer’s instructions. Briefly, cells were harvested and lysed in RIPA buffer, and lysates were clarified by centrifugation at 12,000 × g for 10 minutes at 4°C. Samples, standards, and controls were added to the ELISA plate and incubated with the specific detection antibody at room temperature for the recommended period. Following washes, the substrate solution was added and allowed to develop until a colorimetric change was detectable. Absorbance was measured at 450 nm using a microplate reader. Sample concentrations of 2’3’-cGAMP were calculated by interpolation from the standard curve. All measurements were performed in duplicate.

#### Western blotting

Reserved input lysate or TCA-precipitated materials were analyzed by standard western blotting protocols. Briefly, proteins were separated in Novex 4%–12% acrylamide Bis-Tris gels (NuPage) and transferred to PVDF membranes. Membranes were blocked in Tris-buffered saline containing 0.05% Tween-20 (TBST) and either 5% milk or 3% bovine serum albumin (BSA), according to the antibody manufacturer’s recommendations, and then incubated for 2 h at room temperature with primary antibodies diluted in 1% milk or 3% BSA in TBST with gentle rocking. Blots were washed three times (total 30 min) and incubated with horseradish peroxidase (HRP)-conjugated anti-mouse or anti-rabbit secondary antibodies (Cell Signaling Technologies) for 1 h at room temperature with gentle rocking. Blots were washed again and developed with Pierce ECL Western Blotting substrate (Base, SuperSignal West Pico/Femto, depending on protein levels) and visualized using a Bio-Rad ChemiDoc Imaging System or with HyBlot CL autoradiography films (Denville). Quantification of band intensity was carried out using ImageJ (National Institutes of Health) software.

#### Immunofluorescence

Cells (2 × 10^5^) were seeded in 24-well plates containing 12-mm glass coverslips, and treated with the small molecule drugs as described above. After treatment, cells were fixed with 4% paraformaldehyde in PBS for 20 min or with 100% methanol for 10 min at −20 °C, depending on the antibody specifications, and processed for fluorescence microscopy using standard protocols (https://www.cellsignal.com). Briefly, cells were incubated with primary antibodies of interest diluted in PBS containing 10% FBS and 0.2% saponin for 2 h at room temperature. Cells were then washed three times with PBS and incubated with Alexa Fluor 488- or 568-labeled secondary antibodies (Thermo Fisher) diluted at 1:1000 in PBS/10% FBS/0.2% saponin for 1 h at room temperature. Cells were washed three times in PBS, incubated with DAPI to label nuclei, and mounted with Prolong Gold Antifade Reagent. Imaging was performed using an ApoTome microscope. An average of 10 to 15 stacks of 0.3 μm thickness was acquired.

### Quantification and statistical analysis

#### Puncta number quantification

Puncta were quantified using Imaris 9.3.0 (Bitplane, Zurich). Images were processed using the software “spots” feature with an estimated diameter of 0.5 μm and then corrected using the “different spot sizes: region growing” feature. Thresholds were set using the “Quality” feature and determined manually. The local contrast threshold for region-growing spots was then manually set and maintained. The “surface” feature was used to reconstruct the nucleus of each cell using DAPI staining. Approximately 300–350 proliferating cells and 150–250 senescent cells were analyzed per independent experiment.

#### Colocalization quantification

Colocalization data were processed using Imaris software. In the channels of interest, the appropriate colocalization thresholds were provided in the software “coloc” feature using the “Polygon” tool. The same channel thresholds were maintained for all cells, and the most intense fluorescent overlap was quantified by Imaris as voxels (i.e., overlap between volumetric pixels in either channel). The voxel value for each cell was averaged for each condition and used as a single data point for statistical analysis.

#### Statistical analysis

Figure legends contain all information regarding statistical tests used, number of experimental replicates, dispersion measures used, and specific p-values. Statistical analysis and graph generations was conducted using GraphPad Prism 9 Software. When comparing two datasets, an unpaired two-tailed Student’s t-test was performed. For the comparison of more than two datasets, the one-way ANOVA test was used. A significance level of p<0.05 was considered statistically significant. All data are expressed as Mean±SD from three experimental replicates and are representative of at least two independent experiments. The symbols ∗, ∗∗, ∗∗∗, ∗∗∗∗ indicate significance levels of p<0.01, p<0.05, p<0.01, p<0.001, and p<0.0001, respectively.

## Supplemental Figure Legends

**Supplemental Figure 1:**
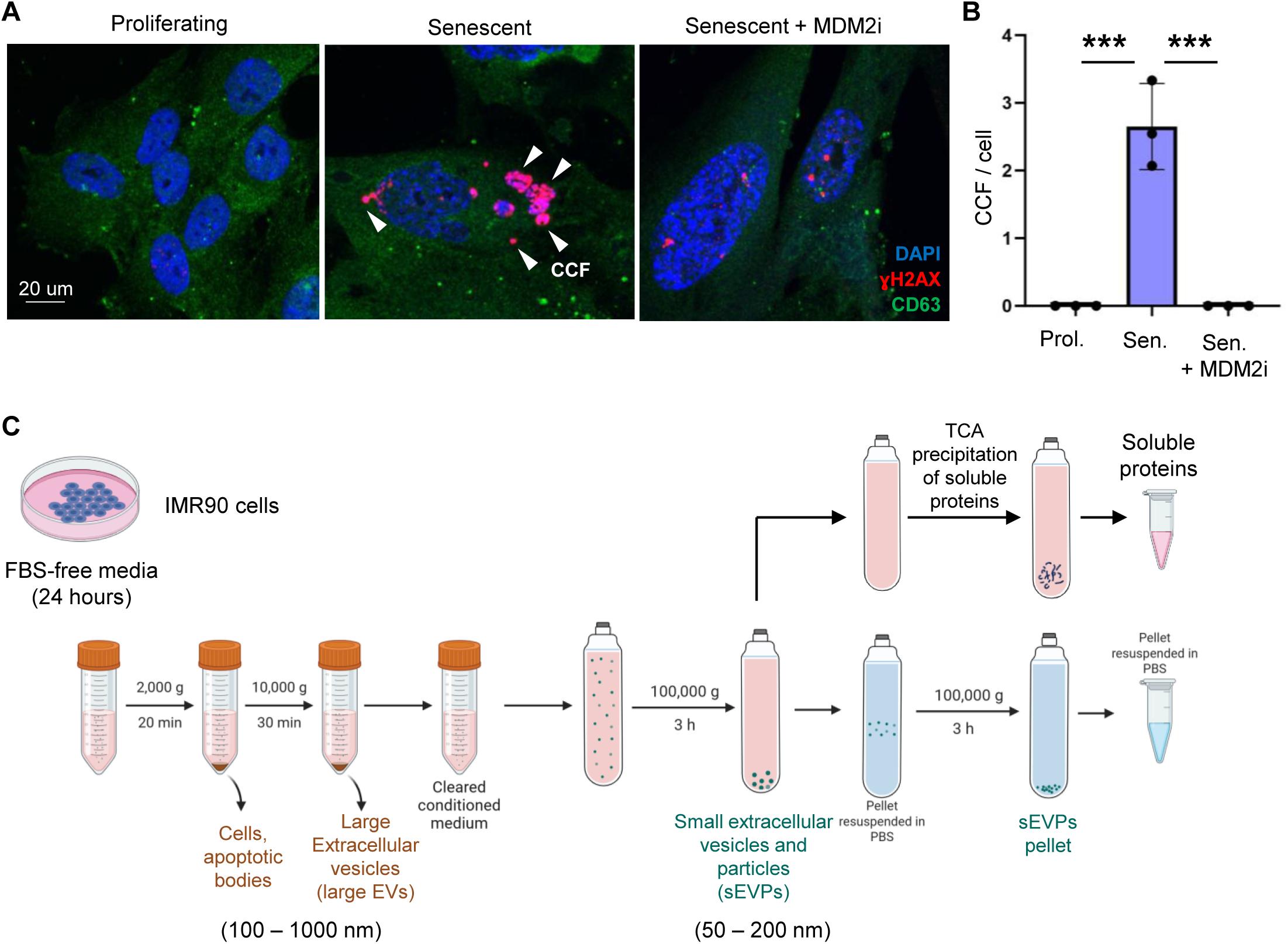
Validation and characterization of CCF-specific secretion. (**A**) Representative immunofluorescence images of proliferating cells treated with DMSO and senescent cells treated with the inhibitor of CCF formation, MDM2 inhibitor RG7388 (MDM2i) or DMSO vehicle, demonstrating decreased CCFs in the treated cells20. (**B**) Quantification of the number of CCFs per cell in proliferating cells treated with DMSO and senescent cells treated with MDM2i or DMSO vehicle. The y-axis represents the total number of CCF, normalized to the total number of nuclei. Data are presented as mean ± SD (n=3 independent experiments). Statistical significance was determined using two-sided one-way ANOVA: ***p=0.0003. (**C**) Schematic representation of centrifugation steps to isolate large extracellular vesicles (EVs), small extracellular vesicles and particles (sEVPs), and soluble proteins. Conditioned media collected from IMR90 cells cultured in FBS-free medium were subjected to differential centrifugation to sequentially remove cells and apoptotic bodies, then large shedding vesicles. The resulting cleared medium was ultracentrifuged at 100,000 × g to pellet crude sEVPs, which were resuspended in PBS and ultracentrifuged again at 100,000 x g to eliminate residual conditioned media and obtain an enriched sEVP fraction. The remaining sEVP-depleted conditioned media were processed by TCA protein precipitation to analyze the soluble protein fraction.

**Supplemental Figure 2:**
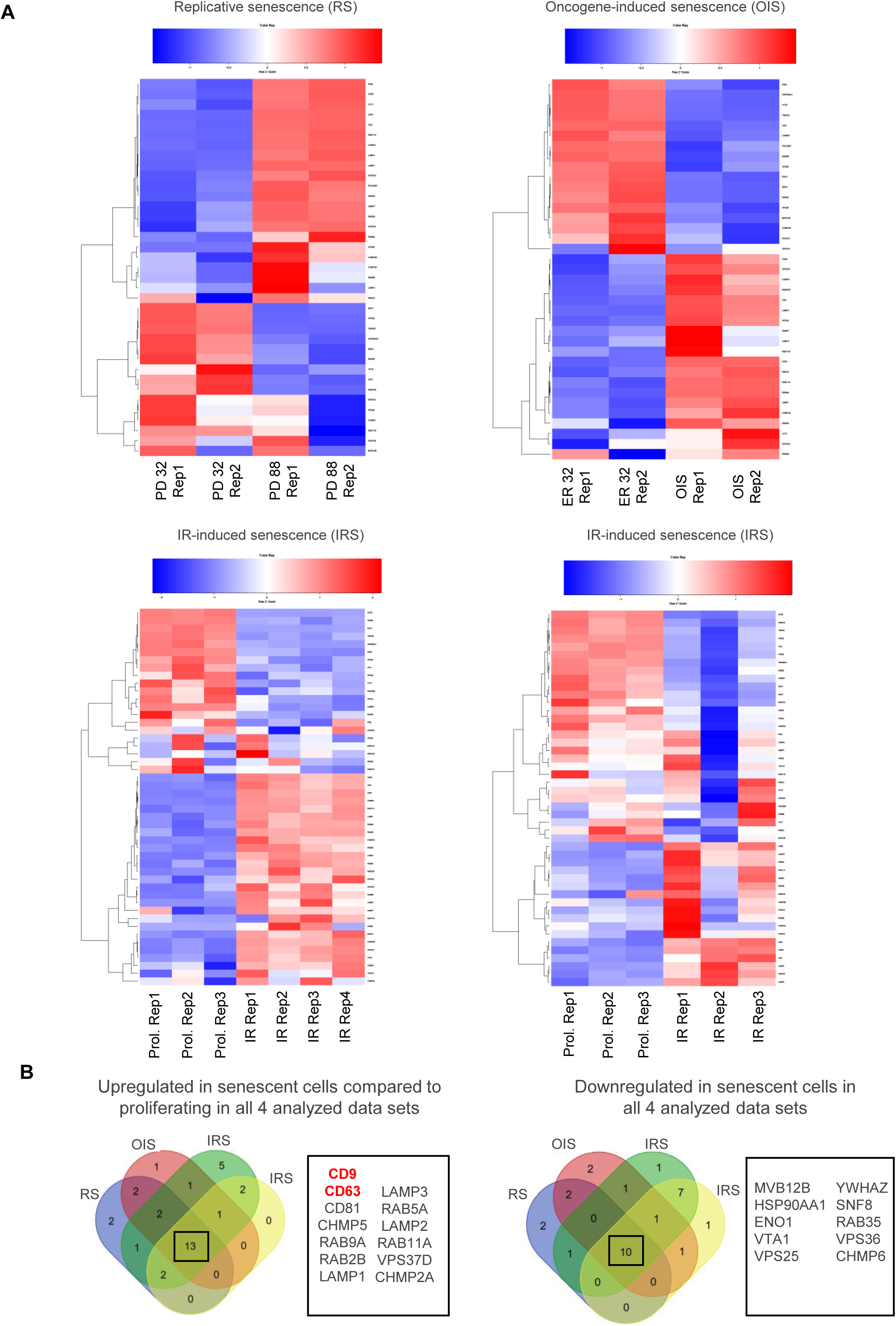
Senescent cells upregulate the set of genes involved in endosomal pathways. (**A**) Analysis of four previously published RNA-seq data sets derived from multiple models of senescence in IMR90 cells, including replicative senescence (RS), oncogene-induced senescence (OIS), and irradiation-induced senescence (IRS)^16,20^, comparing proliferating control versus senescent. Heatmaps display differentially expressed genes associated with the endosomal pathway across the four datasets. In each heatmap, red indicates higher expression and blue indicates lower expression relative to the corresponding proliferating controls. (**B**) Venn diagrams illustrating the overlap of endosomal pathway genes that are consistently upregulated (left) or downregulated (right) across all four RNA-seq data sets corresponding to distinct models of senescence in IMR90 cells (RS, OIS, and IRS). Only genes meeting the criteria for significant differential expression relative to proliferating controls in each dataset were included.

**Supplemental Figure 3:**
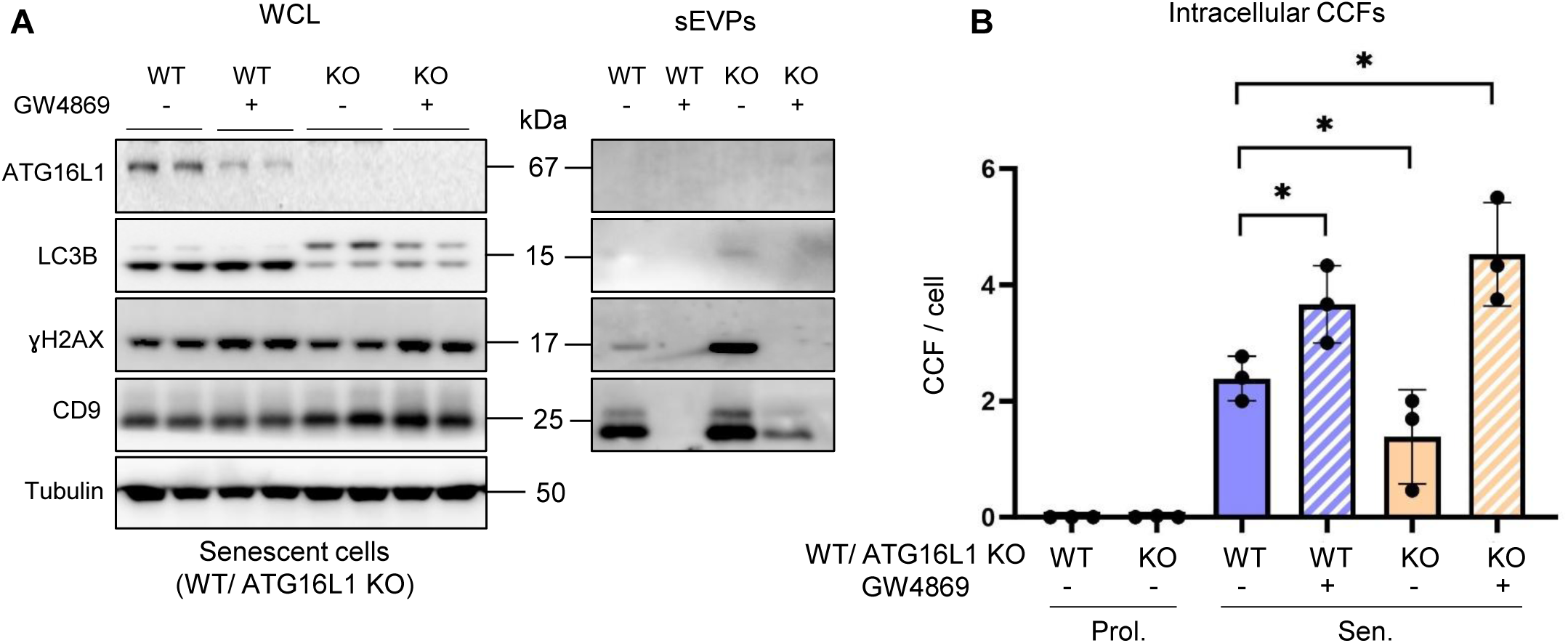
Blocking intracellular degradation of CCFs by ATG16L1 knockout upregulates secretion of the components of CCFs. (**A**) Representative western blot of indicated proteins in whole cell lysates (WCL) and the isolated small extracellular vesicles and particles (sEVP) fractions from conditioned media of senescent wild type (WT) and senescent ATG16L1 knockout (KO) IMR90 cells treated with an inhibitor of the ESCRT-independent pathway, GW4869, and/or DMSO vehicle. GW4869 treatment was applied only during the final 24 h of the conditioning period. WCL samples were normalized to total cell number, and equal amounts of cell lysates were loaded for each condition. The isolated sEVP fractions were resuspended in identical volumes of sample buffer, and equal volumes were loaded per condition. (n=3 independent experiments). (**B**) Quantification of the number of CCFs in proliferating and senescent WT or ATG16L1 knockout IMR90 cells treated with DMSO and/or GW4869. GW4869 treatment was applied only during the final 24 h of the conditioning period. The y-axis represents the total number of CCF, normalized to the total number of nuclei. Data are presented as mean ± SD (n=3 independent experiments). Statistical significance was determined using two-sided one-way ANOVA: *p=0.045, *p=0.032, *p=0.019.

**Supplemental Figure 4:**
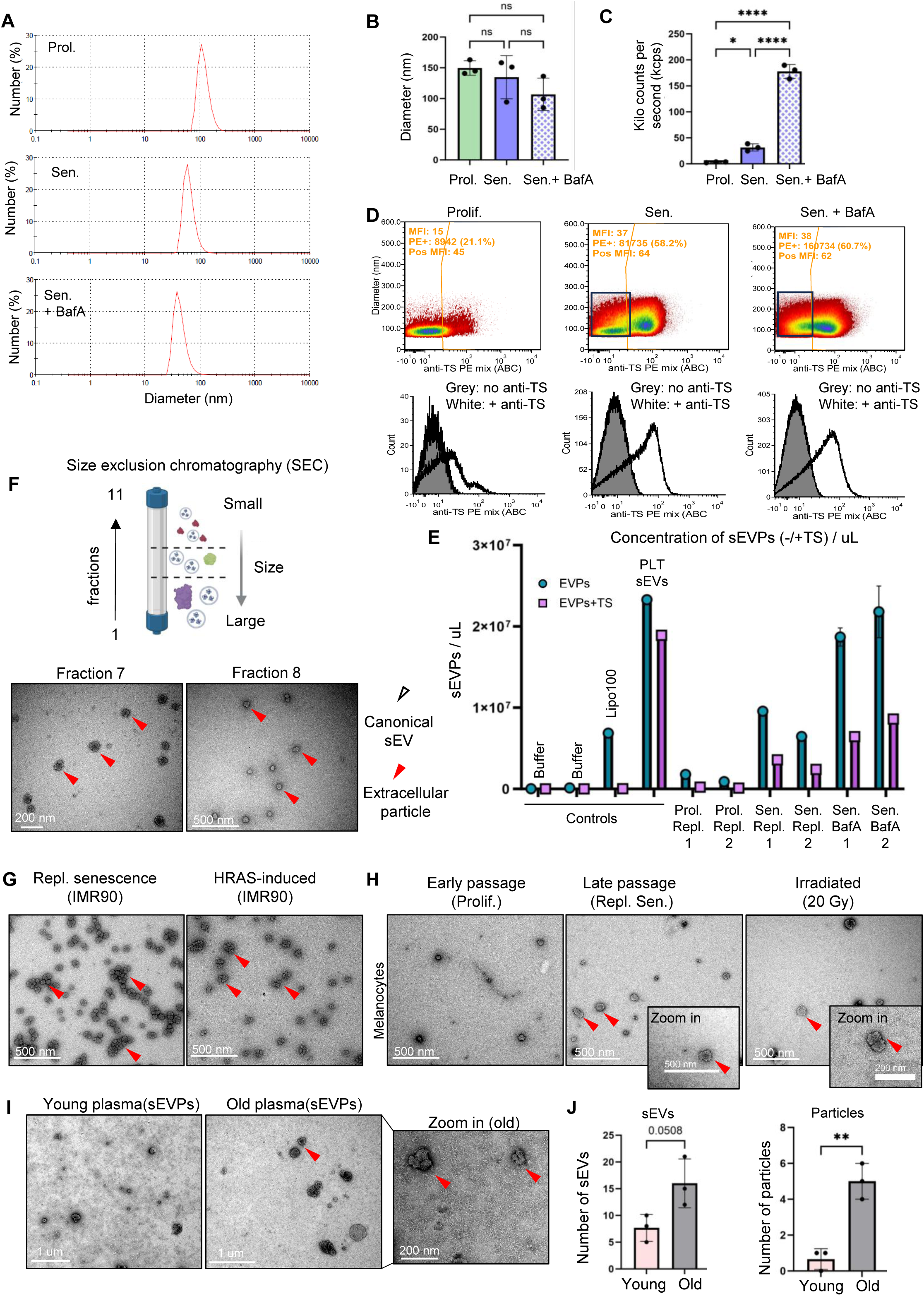
Senescence alters the secretion profile of small extracellular vesicles and extracellular particles across cellular models, perturbation, and aging. (**A**) Representative plots of dynamic light scattering (DLS) analysis of the small extracellular vesicles and particles (sEVP) fractions from proliferating cells treated with DMSO and senescent IMR90 cells treated with lysosomal inhibitor Bafilomycin A (BafA) or DMSO vehicle, showing size distribution (diameter, nm) and number (%), representing the estimated fraction of particle number. BafA treatment was applied only during the final 24 h of the conditioning period. The isolated small extracellular vesicles and particles (sEVP) fractions were resuspended in identical volumes of sample buffer, and equal volumes were loaded per condition. (**B**) Quantification of size distribution determined by DLS of secreted sEVPs from proliferating cells treated with DMSO and senescent IMR90 cells treated with BafA or DMSO vehicle. BafA treatment was applied only during the final 24 h of the conditioning period. The isolated sEVP fractions were resuspended in identical volumes of sample buffer, and equal volumes were loaded per condition. Data are presented as mean ± SD (n=3 independent experiments). Statistical significance was determined using two-sided unpaired one-way ANOVA: ns. (**C**) Quantification of the number of detected sEVPs (kilo count per second) secreted by proliferating cells treated with DMSO and senescent IMR90 cells treated with BafA or DMSO vehicle. BafA treatment was applied only during the final 24 h of the conditioning period. The isolated sEVP fractions were resuspended in identical volumes of sample buffer, and equal volumes were loaded per condition. Data are presented as mean ± SD (n=3 independent experiments). Statistical significance was determined using two-sided one-way ANOVA: *p =0.016, ****p<0.0001. (**D**) Representative histograms of the single vesicle flow cytometry (vFC) analysis of the isolated sEVP fractions secreted by proliferating cells, senescent cells, and senescent cells treated with BafA and/or DMSO vehicle. The sample was stained with the fluorogenic membrane dye vFRed to label and size membrane particles and with a mix of anti-CD9, CD63, and CD81 antibodies conjugated to phycoerythrin (PE) to stain these common tetraspanins. The top panel presents two-parameter histograms of estimated membrane particle diameter versus PE fluorescence, calibrated in units of antibody per cell. The gate for PE positivity was set at the 99^th^ percentile of particle background signal (LOD ∼30 ABC units), and the number and percent of particles exceeding this threshold, as well as their median brightness, are indicated. Events to the right of the vertical gate represent PE-high, tetraspanin-positive particles (canonical sEVs expressing CD9/CD63/CD81), whereas events to the left of the gate represent PE-labeled lipid particles lacking detectable tetraspanins (tetraspanin-negative particles). The bottom panels show the corresponding single-parameter histograms for samples stained with (white) or without antibody (grey). (**E**) Summary of sEVP concentrations for tetraspanin-negative (sEVPs) and tetraspanin-positive (sEVPs+TS) populations within sEVP fractions isolated from conditioned media of proliferating cells treated with DMSO and senescent cells treated with BafA or DMSO vehicle. Buffer-only controls, synthetic liposomes (Lipo100), and platelet-derived sEV standards (PLT sEVs) were included as reference samples. Data represent n = 2 experimental replicates, with two technical replicates per condition (except controls). (**F**) Size exclusion chromatography (SEC) column to separate components of the sEVP fraction based on their size. TEM representative images of the separated sEVP fractions (fractions 7 and 8) secreted by senescent cells using a SEC column, demonstrating pseudoparticles present in fractions 7 and 8 (n=3 independent experiments). (**G**) Representative TEM images of the sEVP fractions isolated from IMR90 cells induced into senescence by replication RS and OIS, demonstrating the presence of extracellular particles. sEVPs were isolated from equivalent numbers of cells for each condition and resuspended in identical buffer volumes prior to downstream analyses (n=3 independent experiments). (**H**) Representative TEM images of the sEVP fractions isolated from proliferating, RS and IRS melanocytes. sEVPs were isolated from equivalent numbers of cells for each condition and resuspended in identical buffer volumes prior to downstream analyses (n=3 independent experiments). (**I**) Representative TEM images of isolated sEVP fractions from the plasma of young (4 months) and old mice (21 months). Equal volumes of plasma were collected from each mouse and pooled within age groups (young vs. old) prior to sEVP isolation. Isolated sEVPs were resuspended in identical buffer volumes before imaging (n = 3 independent experiments). (**J**) Quantification of sEVs and extracellular particles secreted in the plasma of young (4 months) and old (21 months) mice. Equal volumes of plasma were collected from each mouse and pooled within age groups (young vs. old) prior to sEVP isolation. Isolated sEVPs were resuspended in identical buffer volumes before imaging. Data are presented as mean ± SD (n = 3 independent experiments). Statistical significance was determined using a two-sided t-test: **p =0.0029.

**Supplemental Figure 5:**
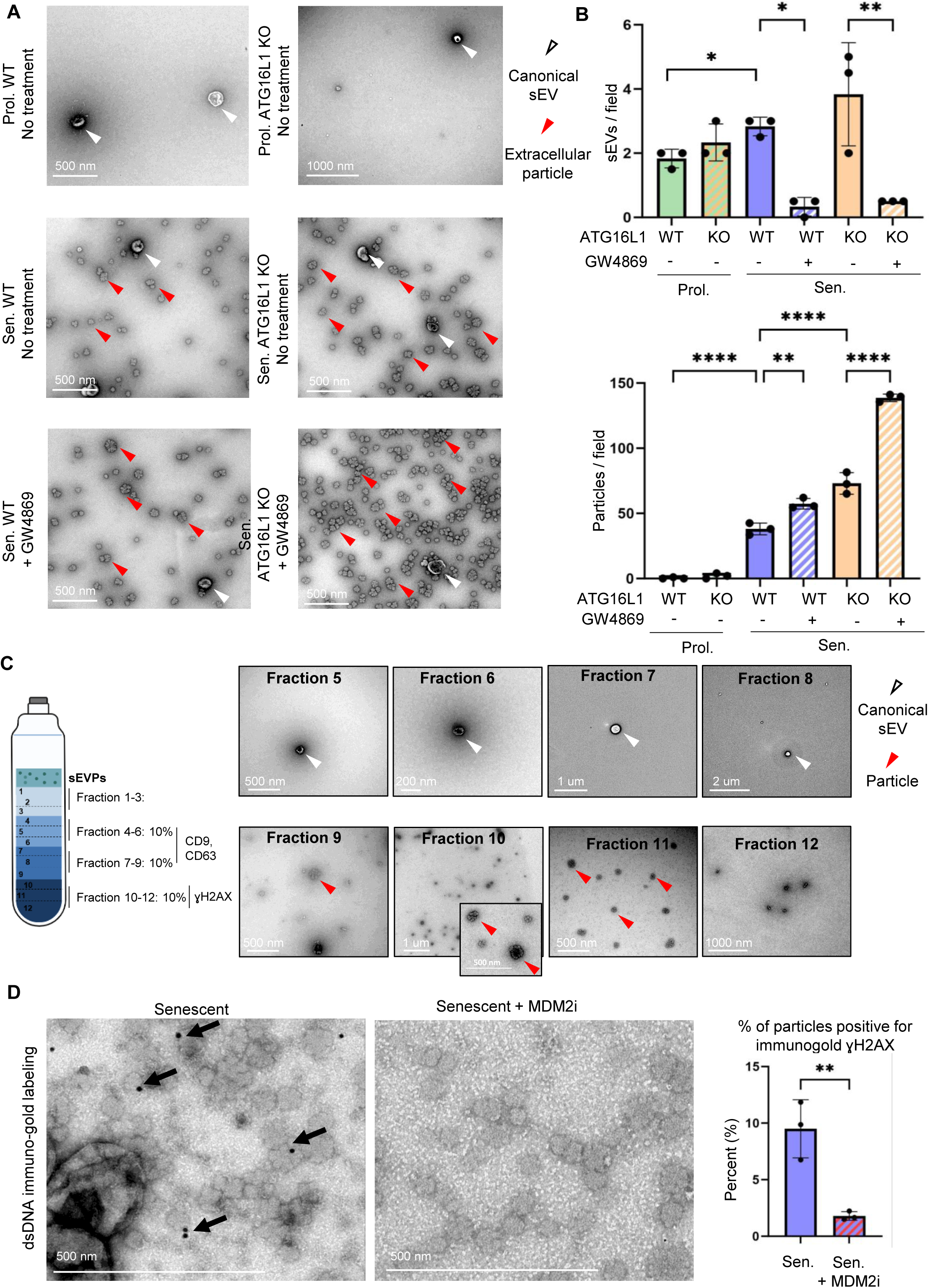
Autophagy inhibition alters the secretion profile of senescent cells, enhancing the release of extracellular particles and associated dsDNA. (**B**) TEM representative images of the isolated small extracellular vesicles and particles (sEVP) fractions secreted by wild-type (WT) proliferating and senescent cells or ATG16L1 knockout (KO) proliferating and senescent cells treated with with an inhibitor of the ESCRT-independent pathway, GW4869 or DMSO. GW4869 treatment was applied only during the final 24 h of the conditioning period. sEVPs were isolated from equivalent numbers of cells for each condition and resuspended in identical buffer volumes prior to imaging. (**C**) Quantification of the isolated sEVs and extracellular particles secreted by WT or ATG16L1 knockout proliferating or senescent cells treated with GW4869 or DMSO vehicle. GW4869 treatment was applied only during the final 24 h of the conditioning period. sEVPs were isolated from equivalent numbers of cells for each condition and resuspended in identical buffer volumes prior to imaging. Data are presented as mean ± SD (n = 3 independent experiments). Statistical significance was determined using two-sided one-way ANOVA: for sEVs/field: * p =0.0132 (control proliferating vs control senescent), 0.0118 (control senescent +/− GW4869 treatment), **p=0.0012 (knockout senescent +/− GW4869 treatment); for extracellular particles/field: **p=0.0018, ****p<0.0001. (**D**) Representative TEM images of different fractions of the separated sEVs and extracellular particles secreted by senescent cells using iodixanol density gradient, demonstrating that fraction 10 contains the majority of the extracellular particles (n = 3 independent experiments). (**E**) Representative TEM images and the corresponding quantification of the dsDNA-immunogold labeling of the isolated sEVP fractions secreted by senescent cells treated with MDM2 inhibitor or DMSO vehicle, demonstrating the presence of dsDNA on extracellular particles secreted by senescent cells. Data are presented as mean ± SD (n = 3 independent experiments). Statistical significance was determined using a two-sided unpaired t-test: **p =0.0068.

